# High specificity meets genomic flexibility in the *Siphamia*-*Photobacterium* symbiosis

**DOI:** 10.64898/2026.03.19.712928

**Authors:** HK Osland, E Neff, A Gaisiner, D Hays, AL Gould

## Abstract

Host-microbe symbioses must balance partner specificity with enough flexibility to remain adaptable to environmental change. The *Siphamia*-*Photobacterium* symbiosis exemplifies this balance, as tropical siphonfish (*Siphamia spp.*) form a highly specific association with the bioluminescent bacterium *Photobacterium mandapamensis*, though it is unknown whether this specificity extends to temperate species of siphonfish. Here, we use long-read genome sequencing and functional assays to characterize the strain-level diversity of symbionts isolated from two temperate siphonfish hosts, *Siphamia cephalotes* and *Siphamia roseigaster*, and compare them to tropical isolates. We found that both hosts exclusively associate with *P. mandapamensis* in their light organs and that temperate strains form host-specific clades despite being collected in close proximity (< 5km). This is consistent with selective host filtering, microhabitat-driven adaptation, or conspecific host seeding. Pangenome analyses showed notable differences in accessory gene content, variation in the *lux*-*rib* operon, and a dramatic expansion of mobile genetic element (MGE) content in a subset of strains. We also identified the first host-derived *P. mandapamensis* isolate that is non-luminescent under laboratory conditions despite an intact *lux*-*rib* operon. Luminescence varied across strains, salinities, and temperature but was not correlated with the presence of *luxF*, with the majority of high-MGE strains exhibiting reduced light output. Together, these results extend the specificity of the *Siphamia*-*Photobacterium* symbiosis into temperate hosts and show that animals can maintain tight symbiont specificity despite the symbiont harboring substantial genomic and phenotypic flexibility at the strain level.

## INTRODUCTION

Microbial symbioses are fundamental to the functioning of marine ecosystems as they shape nutrient cycling, host physiology, and energy flow across diverse environments ^1,2^. The stability of these host-microbe relationships depends on the ability of hosts to balance symbiont specificity with the capacity to adapt to shifting ecological conditions ^3,4^. To maintain this balance, symbiont specificity often reflects how variable the environment is with things such as temperature, salinity, and oxygen, which can result in the divergence of symbionts among closely related microbial strains that are adapted to highly specific niches ^5,6^. Understanding how animal hosts balance fidelity with adaptability is particularly important in the current era of rapid climate change, when shifting conditions can destabilize partnerships between hosts and environmentally acquired microbes ^7–8^.

These varying pressures are reflected at the genomic level, where processes such as gene gain and loss, horizontal gene transfer, and mobile genetic elements (MGEs) can generate plasticity that allow bacteria to adapt to different environments ^9–12^. In symbiotic systems, this plasticity operates against a background of conserved core functions required for colonizing and persisting within desired hosts ^13–15^. Comparing these plastic and conserved regions across closely related strains and hosts makes it possible to pinpoint which genomic features are required for host association and which allow symbiotic bacteria to be ecologically flexible.

Comparative genomics has revealed that substantial functional diversity can exist among strains within a single bacterial species. Closely related strains (≥95% whole-genome ANI) can differ in gene content or regulation in ways that significantly affect host interactions and ecological roles ^16^. With the emergence of advanced sequencing technologies, we can now identify distinct strains and genomic features that are conserved across a species but vary among strains ^17,18^. This is well exemplified by the relationship between *Escherichia coli* and the human microbiome, where closely related strains range from harmless gut residents to virulent pathogens ^19^. Mutualistic systems also exemplify this dynamic as strains vary in their ability to colonize hosts and their effects on the hosts, as seen in the legume-*Rhizobia* ^20^, nematode-*Xenorhabdus nematophila* ^21^, and squid-*Vibrio* systems ^22,23^. These examples illustrate the power of a strain-level approach for studying the genetic mechanisms that govern colonization success and host impact in many symbiotic associations ^24^.

An ideal system for exploring these dynamics is the symbiosis between siphonfish (*Siphamia spp.*) and the bioluminescent bacterium *Photobacterium mandapamensis* ^25^. *Photobacterium mandapamensis* is a subspecies of *Photobacterium leiognathi* in the *Vibrionaceae* family with two circular chromosomes and a broad Indo-Pacific distribution. Its bioluminescence is encoded by the *lux*-*rib* operon, though unlike many *Vibrionaceae*, it is not regulated through quorum sensing ^26–29^. *Photobacterium mandapamensis* can live in a free-living or host-associated state where it has been found to reside in the light organ of multiple fish species ^30^. This dual lifestyle exposes it to ecological and host-mediated selection that makes it a strong model for identifying which genomic features are conserved for symbiosis and which vary with environmental context.

*Photobacterium mandapamensis* has a highly specific relationship with siphonfish, where the symbiont is recruited from the surrounding seawater into a specialized light organ connected to the gastrointestinal tract ^31^. There, the bacteria emit light that the host likely uses for counterillumination during nocturnal foraging ^32–34^. While this host-mediated specificity has been well documented in tropical siphonfish ^35,36^, the symbionts of non-tropical siphonfish are uncharacterized. This gap provides an opportunity to determine whether the high specificity observed in tropical hosts extends to non-tropical siphonfish, and to identify which features of the *Siphamia*-*Photobacterium* mutualism are conserved versus flexible.

In this study, we characterize the symbionts of two temperate siphonfish species that co-occur in the waters surrounding Sydney, NSW, Australia: *Siphamia cephalotes* and *Siphamia roseigaster*. *Siphamia cephalotes* is a temperate species distributed along the southern coast of Australia, where it is primarily found in seagrass- or kelp-dominated habitats in bays and along the open coast ^37^. *Siphamia roseigaster* is primarily a subtropical species, with Sydney marking the southern extent of its range, and is commonly found in sheltered estuaries or bays over sandy or rocky substrates ^38^. *Siphamia roseigaster* is the only siphonfish species that is frequently reported in riverine habitats like the Sydney Harbor estuary, where salinity can drop to brackish or near-fresh conditions (∼5 ppt) after rainfall ^39^.

Investigating symbionts from these two host species provides a broader characterization of temperate symbionts and allows us to compare hosts that differ in biogeographic range and habitat association yet share overlapping ranges. Studying temperate symbionts will also allow us to understand if greater fluctuations in environmental conditions require the symbiosis to be more flexible, and whether the symbionts themselves are more physiologically flexible, as temperate systems experience greater seasonal and thermal variability than tropical reefs ^40^. We address these questions by generating the first whole-genome assemblies of *P. mandapamensis* strains isolated from two temperate siphonfish species and comparing them to previously sequenced tropical symbionts. We use long-read sequencing and functional assays to examine gene content, mobile genetic elements, and performance traits to allow us to distinguish conserved features of the symbiosis from those that vary with host and environment. This work advances our understanding of strain-level specificity in host-microbial relationships and provides information on how host identity and environmental variability shape flexibility within an otherwise highly constrained vertebrate-bacterial symbiosis.

## RESULTS

### Phylogenetic & Pangenome Analysis

We used Oxford Nanopore long-read sequencing to generate the genome sequences of 45 *P. mandapamensis* strains isolated from the light organs of *Siphamia roseigaster* and *Siphamia cephalotes* collected from near Sydney, NSW, Australia (Fig. 1). Assemblies were high quality, with most genomes showing BUSCO completeness scores >99% (Supp. Table 1). A pangenome analysis of the newly sequenced strains, six tropical *P. mandapamensis* strains (host: *S. tubifer*), and three *P. leiognathi* strains (*ljone*.10.1, ATCC25521, *lrivu*.4.1, host: Leiognathidae) included 17,271 total genes across all strains, with 2,638 core genes (≥ 95%), 4089 shell genes (15-95%), 4,241 cloud genes (<15% strains), and 6,303 singletons (single genome). A phylogenetic analysis of the 2,638 core genes revealed that temperate *P. mandapamensis* isolates form distinct clades from tropical strains and cluster into host-specific groups within temperate strains (Fig. 2a). *Siphamia roseigaster* and *S. cephalotes* formed sister clades, with all *S. roseigaster* isolates clustering into a monophyletic group that included a divergent set of strains from a single individual (Sr2) (Fig. 2a). The only exception was strain Sc4.3NL, which was isolated from a *S. cephalotes* host but clustered with *S. roseigaster* isolates. All *P. mandapamensis* strains carried a conserved 14-gene locus encoding glycosyltransferases and regulatory elements that is absent from *P. leiognathi* isolates and homologous to the *syp* locus of *Vibrio fischeri* ^41,42^ (Fig. 2d). *Siphamia cephalotes* strains had a higher number of singletons on average (mean ± s.d. = 105.4 ± 160.3) compared to *S. roseigaster* strains (mean ± s.d. = 41.29 ± 43.08), but the difference between the two groups was not significant (Welch’s t-test: t(17.88) = 1.60, p=0.127) (Supp. Table 2). Within *S. cephalotes* strains, the singleton counts were highly variable (1-534). *Siphamia cephalotes* strains had a higher rate of pangenome accumulation (mean ± s.d. = 1057.07 ± 296.35 clusters/genome) compared to both *S. roseigaster* (mean ± s.d. = 340.65 ± 103.99 clusters/genome) and *S. tubifer* strains (mean ± s.d. = 405.01 ± 41.77 clusters/genome), indicating a more ‘open’ pangenome in *S. cephalotes* symbionts (Fig. 2b; permutation test on accumulation slopes: Sc > Sr, p = 0.03, Sc > StP, p = 0.02).

**Figure 1.**
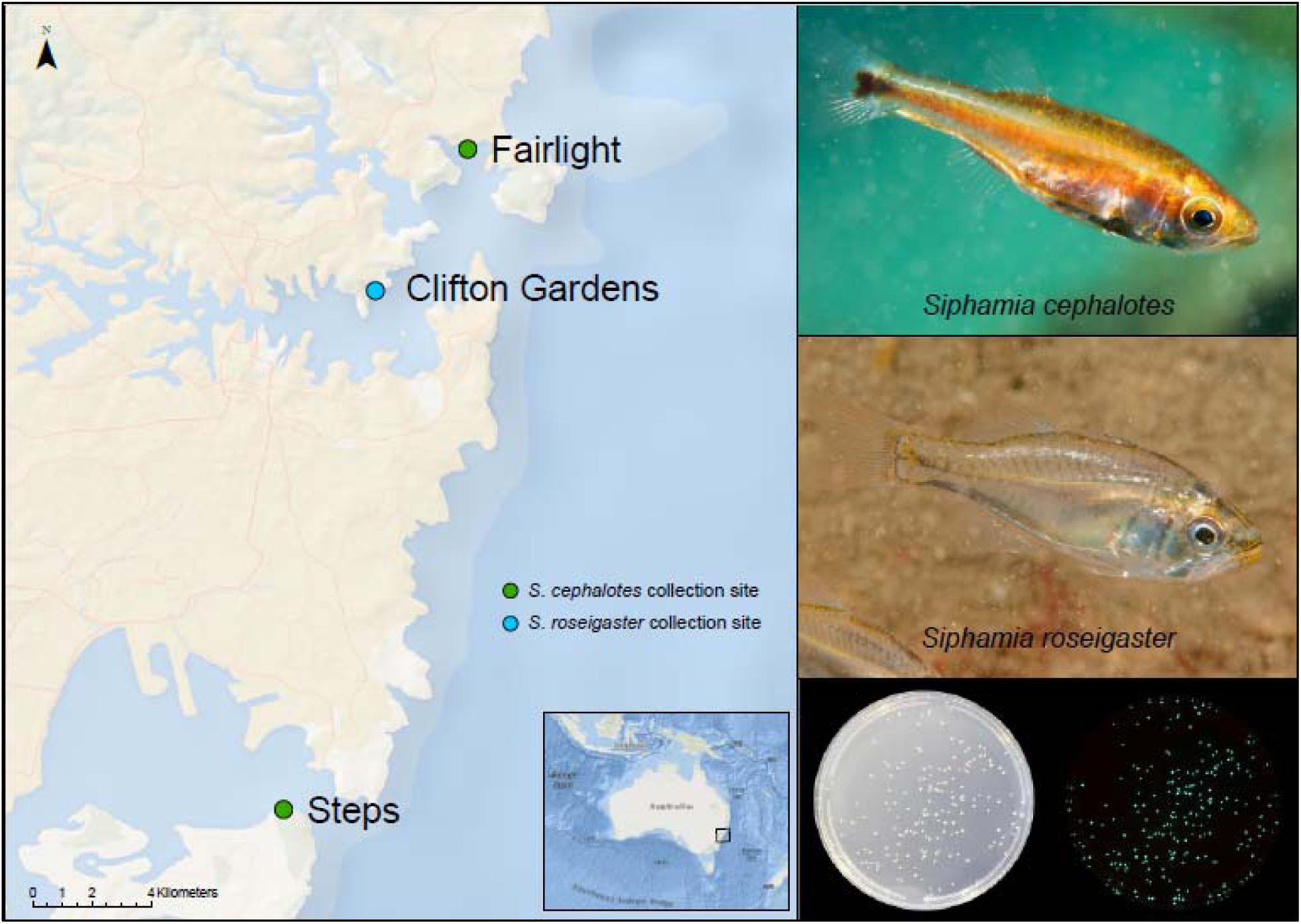
Collection sites, host fishes, and *P. mandapamensis* colonies. (A) Map of siphonfish collection sites in New South Wales, Australia, showing the three locations where *S. cephalotes* and *S. roseigaster* were sampled. (B) Images of adult *S. cephalotes* and *S. roseigaster* showing differences in body coloration by Erik Schlögl (iNaturalist, CC BY-NC 4.0) (C) Bioluminescent *Photobacterium mandapamensis* colonies isolated from *S. tubifer* light organs in ambient and dark light conditions.

**Figure 2.**
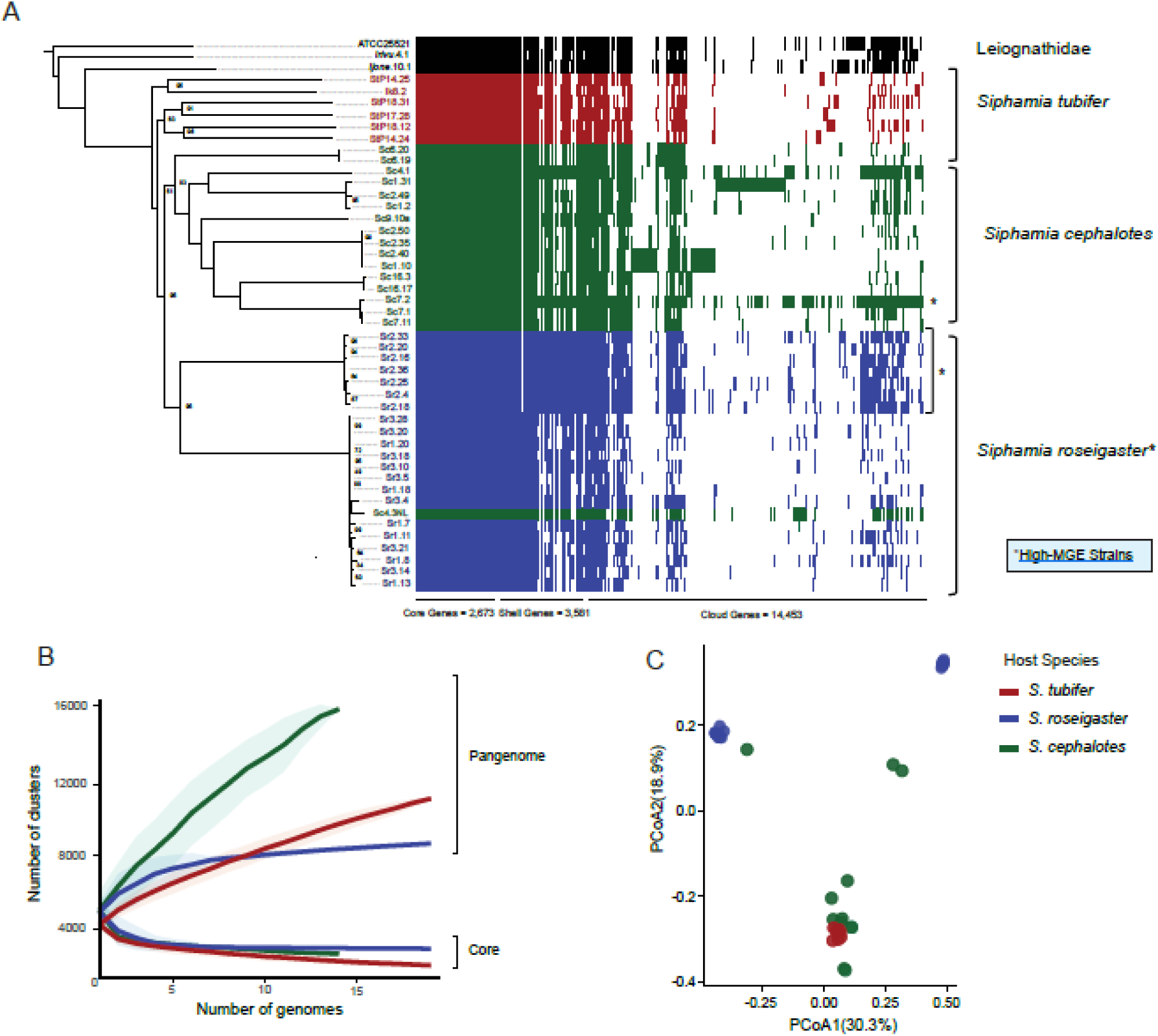
Core-genome phylogeny and pangenome analysis of *P. mandapamensis* strains. (A) Maximum likelihood phylogeny and pangenome analysis of *P. mandapamensis* strains using 2,673 core genes, 1000 bootstraps, and IQ-TREE model GTR+F+I+G4. Labels are colored by host species (*S. tubifer*, *S. roseigaster*, *S. cephalotes*, and *P. leiognathi* reference strains) with bootstrap support values listed if below 95%. Phandango plot of gene-presence absence with singletons removed. Asterisks represent high-MGE strains (B) Core- and pangenome accumulation curves based on the Roary gene-presence-absence matrix. Ribbons represent ± 1 SD. (C) PCoA plot using Jaccard distances computed from accessory genomes colored by host-association.

### Variability in the lux-rib operon and bioluminescence

The pangenome analysis identified notable differences in gene content between temperate and tropical isolates, as well as host-associated divergence within temperate strains (Fig. 2). One region of variation was the *lux-rib* operon that is responsible for light production (Fig. 3a). Previously sequenced *P. mandapamensis* isolates from tropical siphonfish carry the full operon, which consists of the core luciferase genes *luxCDABFE*, upstream riboflavin biosynthesis genes (*rib*), and the downstream accessory genes *lumP* and *lumQ*, which modulate light intensity and spectral output ^28,43–45^. In contrast, temperate isolates showed structural variation in the *lux*-*rib* operon. Only a single temperate strain (Sc4.3NL) carried the full *luxF* gene, an accessory component shown to enhance light output by reducing inhibitory byproducts of the luciferase reactions, while all other temperate strains lacked *luxF* or carried partial deletions (78- 749 bp; Fig. 3a) ^46^. The presence of *luxF* in *S. tubifer*-associated strains and its partial presence in *S. roseigaster* isolates coincided with additional copies of *luxC* and *luxD*, which were absent from all other *S. roseigaster* and *S. cephalotes* strains. These additional copies were located outside the primary *lux*-*rib* operon in another region of chromosome two. Across temperate isolates, the absence of *luxF* was consistently associated with partial deletions in *lumP* (317-321 bp; Fig. 3a).

**Figure 3.**
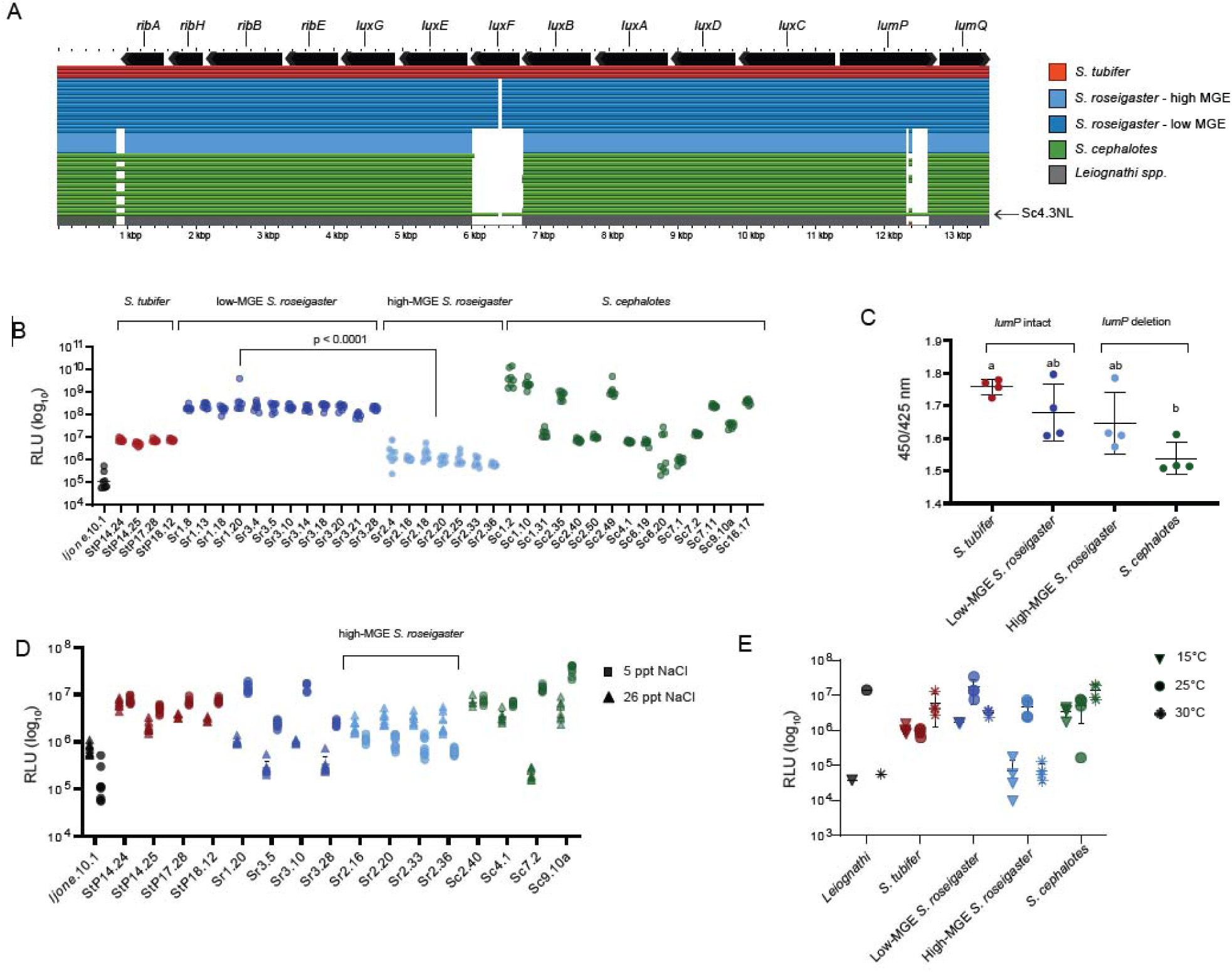
Lux-operon and luminescence across *P. mandapamensis* strains. (A) Comparative organization of the *lux*-*rib* operon across *P. mandapamensis* genomes illustrating host-associated presence, absence, and truncation of certain genes. Each row represents a strain with coloring according to host species, with arrows showing individual genes and their orientation. (B) Luminescence of *P. mandapamensis* strains defined as the maximum RLU value obtained during the 24-hour growth period. Within *S. roseigaster*, low-MGE strains were significantly more luminous than high-MGE strains (p < 0.0001). Full pairwise comparisons among host groups are provided in Supplemental Table 4. (C) Host-associated shifts in emission color using 450/525 nm ratio, where higher values indicate relatively bluer emission and lower values indicate relatively greener emission (D) Luminescence values of *P. mandapamensis* under standard (26 ppt) and low-salinity (5 ppt) conditions and (E) varying temperatures (15°C, 25°C, 30°C).

To determine if there was a functional consequence to this variation in the *lux*-*rib* operon, we measured the luminescence of strains at varying temperatures (15°C, 25°C, 30°C) and salinities (5 ppt, 26 ppt). We found variability that corresponded with host-association, MGE content, salinity, and temperature (Fig. 3b-e). Under standard conditions (25°C, 26 ppt NaCl), *S. roseigaster*-derived strains had luminescence levels comparable to tropical *S. tubifer*-derived strains. In contrast, *S. cephalotes* strains were more variable, with some exceeding tropical levels and others falling below. Under standard conditions, high-MGE strains (originating from host Sr2) had lower luminescence than strains from other *S. roseigaster* hosts (p < 0.0008). The majority of strains had lower luminescence under low-salinity conditions than under standard conditions (p < 0.001; F(1,182) = 561.4), except for high-MGE Sr2 strains, which produced higher luminescence under low-salinity conditions than under standard salinity (p < 0.0001; F(1,56) = 84.14). In contrast, the high-MGE *S. cephalotes* strain (Sc7.2) did not show reduced luminescence relative to other *S. cephalotes* isolates and had a higher luminescence in standard salinity compared to low salinity.

Luminescence was temperature-dependent (p < 0.0001; F (2,381) = 366.27) and differed among host-associated groups (p = 0.0059; F(4,12) = 6.24). Across temperatures, *S. tubifer* and *S. cephalotes* strains maintained relatively high luminescence values, whereas high-MGE S. roseigaster-associated strains showed the largest temperature-dependent shifts, with the highest luminescence at 25°C compared to 15°C or 30°C. A linear model of log_10_-transformed luminescence values with salinity, IS copy number, temperature, and host as predictors explained roughly half of the variation (R² = 0.49). In a Type II test, IS copy number, temperature, and host species were significant (all p < 10^-14^), while salinity was not significant after controlling for the other predictors (p > 0.05). Given the differences in the presence of *lumP*, we also measured the emission properties between strains (450/525 nm ratio) (Fig. 3c). *Siphamia tubifer* strains, which had an intact *lumP* gene, had light emission shifted toward bluer wavelengths (mean ± s.d. = 1.76 ± 0.02), and *S. cephalotes* strains that carried partial *lumP* deletions had light emission shifted toward greener wavelengths (mean ± s.d. = 1.54 ± 0.05). Both low-MGE and high-MGE *S. roseigaster* strains showed intermediate values (mean ± s.d. = 1.66 ± 0.09). One *S. cephalotes* strain (Sc4.3NL) was non-luminescent under all conditions tested, including assays with decanal supplementation. When we grew Sc4.3NL in co-culture with a luminous strain (Sr3.5), the luminescence of the co-culture strain was reduced compared to monoculture controls (Supp. Fig 3). To our knowledge, Sc4.3NL represents the first “dark” *P. mandapamensis* strain isolated from a siphonfish light organ in the wild. Overall, these findings show that the *lux*-*rib* operon varies between temperate and tropical *P. mandapamensis* and among temperate strains from different hosts, suggesting that luminescence and regulation may shift across geography and host.

To determine if there is a trade-off between light production and growth rate, we also measured growth rates under standard and variable conditions and found no significant correlation between luminescence and growth rate (*r_s_*(15) = 0.17, p = 0.52). All strains showed their highest growth rate at 26 ppt NaCl, with significantly reduced growth at lower salinities (Supp. Fig 1). Growth rate was significantly lower at 15°C compared to the other temperatures for all strains (p < 0.0001). Growth rate did not differ significantly by host association (p > 0.8) or MGE content.

### Mobile genetic element variation

Gene content comparisons revealed substantial variation in the abundance of mobile genetic elements (MGEs) among *P. mandapamensis* strains. In contrast to previously sequenced *P. mandapamensis* genomes, which contain few to no insertion sequences (0-15 IS elements) ^29,47^, seven distinct strains isolated from a single *S. roseigaster* individual (Sr2) had markedly elevated IS content, with 452-470 ISs per genome (mean ± s.d. = 11.41% ± 0.20 of the genome; Fig. 4A). These high-MGE Sr2 strains formed a distinct subclade within the *S. roseigaster* clade in the core-genome phylogeny (Fig. 2). One *S. cephalotes* isolate (Sc7.2) also showed elevated IS content, with 299 IS elements (8.20% of the genome). The ISs from these high-MGE strains belonged to 11 IS families (dominated by IS5, IS66, and IS630), with the number of IS copies from each family highly comparable across the eight strains (Fig. 4; Supp. Table 3). The IS elements were distributed across both chromosomes, with a higher density on chromosome 2 (mean ± s.d. = 59.52% ± 5.43) than on chromosome 1 (mean ± s.d. = 40.48% ± 5.43). Sr2 strains shared many common insertion sites but also carried strain-specific insertions, while Sc7.2 had a more distinct IS landscape (Fig. 4d). Phylogenetic analysis of IS sequences showed that *P. mandapamensis* ISs clustered primarily with ISs from other *Vibrionaceae* (Fig. 4c; Supp. Figs 4-8). Kimura distance distributions showed a small number of older copies with two major expansion events (Fig. 4e). Beyond single IS elements, MobileGeneticElement finder identified between 150-200 putative composite transposons in each Sr2 genome and 95 in Sc7.2 (Supp. Table 3).

**Figure 4.**
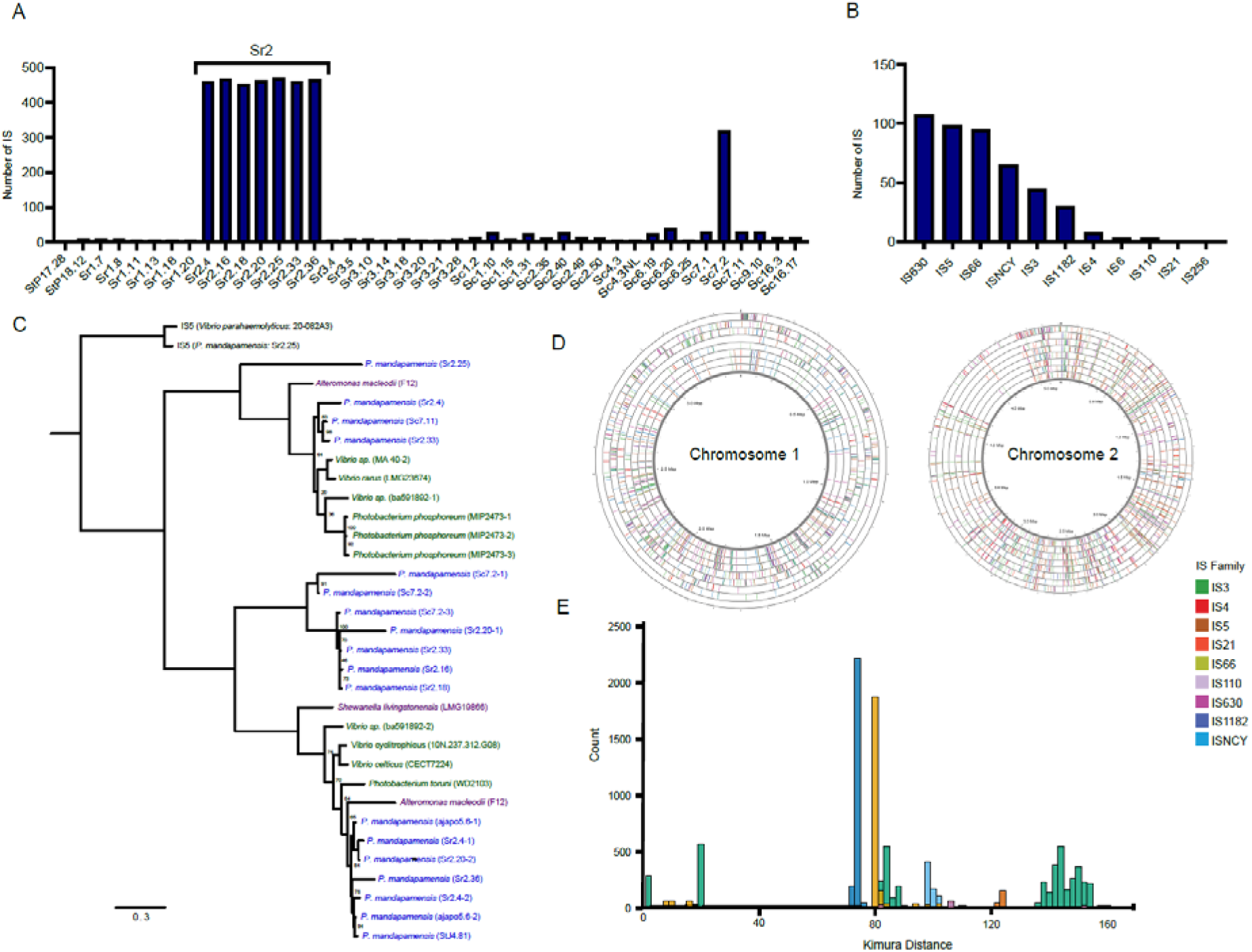
Mobile genetic elements and insertion sequences of P. mandapamensis. (A) Number of insertion sequences per genome. (B) Composition of IS families in the high-MGE strain Sr2.20 based on ISEScan annotations. (C) Phylogenetic tree for IS6 family detected in *P. mandapamensis* showing that most IS copies cluster within *Vibrionaceae*-associated lineages. Bootstrap support values listed if below 95%. (D) Circular chromosome plots of high-MGE strains plotted on a consensus genome showing the genomic locations and density of IS insertions. Strain order from inner to outer ring: Sr2.16, Sr2.20, Sr2.33, Sr2.25, Sr2.4, Sr2.18, Sr2.36, Sc7.2. (E) Kimura distance distributions for IS copies within high-MGE strain Sr2.20 that illustrate the existence of resident elements and two more recent expansion events.

### Accessory gene variation

We identified several accessory genes that varied among strains based on host association. Low-MGE *S. roseigaster* strains carried a contiguous block of 296 genes that were present in > 88% low-MGE *S. roseigaster* strains but only < 10% of other strains. More than half of these genes were annotated as hypothetical proteins (191/296), while the remaining genes were primarily annotated as phage/superinfection exclusion, MSHA/flagellar functions, transporters, and regulatory proteins (Supp. Data 2). Oxidative stress-associated genes (*gstB*, *selB*, *mdtI*), metabolic genes (*galK*), and a T3SS-associated gene, *sctC*, were present in S. cephalotes and S. tubifer strains but absent from *S. roseigaster* strains ^43^.

## DISCUSSION

Here, we generate the first whole-genome assemblies of temperate *P. mandapamensis* strains isolated from siphonfish light organs and show that host specificity is conserved in temperate habitats. We found that symbionts from *S. roseigaster* and *S. cephalotes* form distinct host-associated clades that are separate from tropical lineages and exhibit substantial genomic flexibility. This combination extends previous observations of high specificity in the *Siphamia-Photobacterium* symbiosis and reveals unexpected levels of strain-level plasticity in the symbionts.

Our genomic analysis of the symbiont revealed strong host-associated structuring despite minimal geographic separation. Sampling sites were located less than 5 km apart within the same bay (Fig.1), though the two species typically segregate by habitat type. *Siphamia roseigaster* is commonly found over sandy and rocky substrates, while *S. cephalotes* is more commonly found in seagrass or kelp beds ^37,38^. The *S. roseigaster* site is located slightly deeper into the Sydney Harbor Estuary between the two *S. cephalotes* sites, yet *S. cephalotes* symbionts were more similar to one another than to those from *S. roseigaster*. This indicates that host identity is more strongly correlated with strain identity than geographic location. Such fine-scale structuring could arise from highly specific host filtering among closely related bacterial strains (>95% whole-genome ANI), ecological partitioning of free-living *P. mandapamensis* across microhabitats, or host enrichment by conspecific adults that seed local environments with resident strains. Under the latter two scenarios, larvae would encounter *P. mandapamensis* in patchy, locally distinct microenvironments, where colonization depends on when and where larvae intersect strain-enriched pockets. In any of these scenarios, this symbiosis exemplifies a high degree of specificity at the subspecies level between siphonfish and *P. mandapamensis*. One candidate mechanism that may contribute to this high specificity is a conserved 14-gene polysaccharide biosynthesis and export locus that includes glycosyltransferases, an *etk*-like tyrosine kinase, and *syp*-homologous regulator genes. Because this locus is homologous to the *V. fischeri syp* region and absent in *P. leiognathi* ^42^, it may influence host-associated aggregation or attachment behaviors that contribute to specificity between siphonfish and *P. mandapamensis*. As our sampling represents a snapshot in time of a limited number of hosts and sites, additional sampling across seasons and years will be needed to determine how stable this partnership is over time and through recruitment events.

A pangenome accumulation analysis showed that *S. cephalotes* strains have a more “open” pangenome than strains from *S. roseigaster* and *S. tubifer*, which may relate to the thermal distributions of the hosts. *Siphamia cephalotes* is the only host in this study with a fully temperate range, whereas *S. roseigaster* is primarily sub-tropical with a small temperate range, and *S. tubifer* is sub-tropical and tropical. *Siphamia cephalotes* populations also vary more seasonally compared to the other two hosts, as they become less abundant during winter months in certain areas of their range ^49,50^. This seasonal host limitation may reduce opportunities for *P. mandapamensis* to reliably encounter their hosts and increase the need to persist longer in the environment. Another factor could be the greater abiotic variability of temperate habitats relative to tropical habitats, where stronger seasonal fluctuations may select for more open genomes ^51^. This more facultative lifestyle could favor the retention of a larger, more flexible accessory genome, in contrast to the more streamlined genomes expected for tropical symbionts that encounter more stable conditions and consistently available hosts ^52,53^.

Despite the persistence of strain-level specificity within the *Siphamia*-*Photobacterium* symbiosis in temperate habitats, *P. mandapamensis* strains showed significant differences in genome content. These differences illustrate the remarkable genomic plasticity that microbial symbionts can exhibit, even within highly specific partnerships. Notably, this plasticity extended to the *lux*-*rib* operon, where strains from different hosts maintained the same core bioluminescence machinery, but varied in accessory genes responsible for optical tuning. These changes in accessory genes related to luminescence were further structured within temperate isolates, indicating that bioluminescence phenotypes can be tuned to host and environmental conditions, with different brightness or spectral profiles favored under varying conditions.

Genomic differences in the *lux*-*rib* operon were associated with varying patterns of luminescence output that varied with host association, IS copy number, salinity, and temperature. Luminescence corresponded with host association, which is consistent with host-linked differences among symbiont lineages. Under standard conditions (25°C, 26 ppt), *S. roseigaster* and *S. tubifer*-derived strains showed relatively uniform light production, while *S. cephalotes* symbionts spanned a wide range of luminescence levels. This potential to be brighter than tropical strains was unexpected, given that *S. cephalotes* strains lack *luxF*, a flavin-cleaning protein typically associated with enhanced luminescence ^54^. This suggests that regulatory differences or broader genomic context play a role in determining light output. Historically, *luxF* has been used as a distinguishing feature between *P. leiognathi* and *P. mandapamensis* ^55^, however, our results show that *luxF* is absent in multiple *P. mandapamensis* lineages and is no longer a reliable taxonomic marker. High variability in luminescence among *S. cephalotes* strains may be related to their more open genome, as greater genomic flexibility could broaden physiological performance and result in more diverse phenotypes.

High IS copy number also had a significant effect on luminescence in some strains, as high-MGE strains from Sr2 were significantly less bright than their low-MGE Sr counterparts. There were no insertions that directly interfered with luminescence-related genes, but a high IS copy number may affect metabolism, regulation, or stress responses that indirectly impact light production. However, the high IS copy number in Sc7.2 did not significantly affect luminescence, suggesting that other factors in the Sr2 strains are responsible for the reduced luminescence.

Temperature also affected luminescence, with higher temperatures yielding higher maximum luminescence values, except in *S. roseigaster* strains, where symbionts peaked at 25°C. This provides further support for host-associated differences in thermal sensitivity. Salinity stress shaped luminescence phenotypes but contributed comparatively less variation once other predictors were included. The most notable result from this experiment was the high performance of high-MGE *S. roseigaster* strains under low-salinity conditions. While most strains had reduced light output under low-salinity conditions, a subset of high-MGE strains showed enhanced luminescence at lower salinities, indicating that salinity tolerance and luminescence can diverge among closely related strains. This pattern suggests that environmental history may affect luminescence, consistent with the possibility that the Sr2 strains originated in brackish or river-influenced habitats where selection favors luminescence under low-salinity conditions. This interpretation aligns with the ecology of *S. roseigaster*, as it is the only siphonfish species documented in brackish waters. Given the location of fish collection, this may indicate that Sr2 acquired high-MGE strains in the deeper riverine areas of the Sydney River system, before settling closer to the coast in an area of standard ocean salinity (Fig. 1). As we completed our assays under controlled laboratory conditions, we will need to complete future experiments in free-living or host-associated bacteria to determine if these patterns correspond with performance in the wild.

There was an additional link between *lux*-*rib* gene content and luminescence output, though the pattern was not applicable across all host groups. Strains with an intact *lumP* (*S. tubifer*) generally shifted light emission toward bluer wavelengths rather than green, which is consistent with prior work suggesting that *lumP* tunes emission towards shorter, bluer wavelengths. This adaptation may be advantageous in clear tropical waters where violet/blue wavelengths are transmitted more efficiently, and the downwelling light field is blue-weighted ^56,57^. Selection in blue-weighted tropical or clearer waters may favor their retention in *S. tubifer* strains, improving spectral matching for counterillumination, while this advantage is reduced or absent in temperate or more turbid environments of *S. cephalotes* strains.

We also identified the first naturally occurring *P. mandapamensis* strain (Sc4.3NL) isolated from a siphonfish light organ that produced no detectable light under laboratory conditions despite harboring an intact *lux*-*rib* operon (Fig. 3). This strain clustered phylogenetically within the *S. roseigaster* clade, even though it was isolated from an *S. cephalotes* host, and remained dark under all laboratory conditions, including assays with decanal supplementation and partially spent medium of other luminous strains (Supp. Fig 4). Although we cannot exclude the possibility that Sc4.3NL is luminescent under non-laboratory conditions due to a lack of host-associated or environmental cues, its existence raises questions about host enforcement and the potential for non-contributing symbionts in this system. In other mutualisms, such as the legume-*Rhizobia* and squid-*Vibrio* systems, host control mechanisms can identify and expel “cheater” strains that prevent the collapse of the partnership ^58–61^. Whether siphonfish can similarly detect or sanction non-luminous *P. mandapamensis* is unknown, but testing the competitive fitness of Sc4.3NL in mixed infections, its persistence within light organs, and host physiological responses will provide an avenue for probing enforcement mechanisms in this symbiosis. Overall, these patterns suggest that even within a highly specific symbiosis, the *lux*-*rib* operon and luminescence of *P. mandapamensis* are flexible and can be remodeled across environments and host species without disrupting partner specificity. These results also show that luminescence is shaped by a combination of gene content, environmental history, and host ecology, rather than by the presence or absence of a single gene.

Another form of genomic flexibility in *P. mandapamensis* is the extreme expansion of mobile genetic elements we observed in a subset of strains that was in contrast with previously sequenced *P. mandapamensis* genomes that contain very few insertion sequences. These are, to our knowledge, the highest counts reported for any member of the *Vibrionaceae* family and align more closely with counts typically reported in microbes from extreme or unstable environments ^62–64^. Most of these IS copies were phylogenetically nested within IS from other *Vibrionaceae* lineages, suggesting expansion of resident elements rather than acquisition from distant bacteria. The similar IS family ratios across strains further support an ancestral IS repertoire that was retained during expansion. IS elements were distributed across both chromosomes, with Sr2 strains sharing many insertion sites, while Sc7.2 had a distinct insertion sequence landscape that points to a more isolated expansion. Roughly half of the IS copies in high-MGE strains were classified as putative composite transposons, suggesting a potential capacity to mobilize cargo genes that could serve as vehicles for adaptive traits ^65^. Inspection of the genes carried within these elements showed that most encoded proteins of unknown function and did not reveal clear enrichment for any specific functional categories, though some are strong composite transposon candidates due to flanking IS elements and cargo gene functions (Supp. Table 5).

The expansion of IS elements in this subset of strains could have occurred during the free-living stage of *P. mandapamensis* or within the siphonfish host. In the environment, exposure to recurrent DNA-damaging stressors like UV radiation, oxidative damage, and exposure to pollutants can activate the SOS response and lead to increasing excision and transposition rates ^66–71^. This is consistent with experimental work that has shown that resident ISs become more active under stress, with IS-mediated mutations contributing disproportionately to adaptation during episodes of nutrient limitation, oxidative stress, high temperatures, and antibiotic exposure ^70–73^. Under this scenario, *P. mandapamensis* in fluctuating coastal and riverine habitats could experience localized bursts of SOS activation and IS expansion, generating the high-copy-number genotypes we observe.

Siphonfish are receptive to symbionts 7 days post-release from the paternal mouth and stay receptive for the duration of their ∼4-week pelagic larval stage, so fish settling into similar habitats may still recruit strains with different stress histories depending on when and where colonization occurs. This life history offers an explanation for why Sr2 and Sc7 carried high-MGE strains, whereas conspecific fish collected at the same sites did not. If high-MGE strains did arise in response to environmental stress, Sr2 and Sc7 highlight how fine-scale differences in coastal conditions can drive dramatic genomic change in bacteria that have phenotypic consequences on the host bacterium.

Alternatively, MGE proliferation could have occurred post-colonization within the host light organ, where host-associated conditions, population bottlenecks, or chronic physiological stress could have led to the expansion ISs ^67,74–76^. However, only one of the seven sequenced symbionts within Sc7 carried a high number of MGEs, which argues against a uniform, deterministic host effect. Another relevant pattern is that large IS expansions are frequently associated with the onset of genome reduction in obligate symbionts ^77–80^. That trajectory seems less likely here because *P. mandapamensis* remains a facultative, environmentally acquired partner, and most of the siphonfish we collected had a low number of MGEs. Whether these expansions originated in the environment or inside the host are both compelling, as they either point to strong environmental structuring of symbiont genotypes or to the light organ itself as a hotspot of microbial genome evolution ^81^.

Elevated IS counts of this magnitude are notable because IS and composite transposons can have functional consequences on microbes that can accelerate adaptation to changing environments or cause deleterious insertions into functional genes ^71,73,75,82,83^. MGE activity can generate phenotypic diversity and promote local adaptation on short timescales, but it can also impose significant fitness costs by destabilizing genomes and disrupting essential regulatory or metabolic genes ^75,84,85^. Our data may reflect these potential trade-offs, as high-MGE strains from the Sr2 fish had lower luminescence levels under standard conditions but became relatively brighter under low-salinity conditions. This pattern may not be driven by MGEs, as the single *S. cephalotes* high-MGE strain (Sc7.2) did not see similar negative effects, but could reflect local adaptation of symbionts associated with *S. roseigaster*, as it is the only siphonfish species frequently reported from riverine habitats where salinity can be decreased. In this context, elevated MGE loads could be double-edged, as they may have allowed *P. mandapamensis* to adapt to low-salinity environments, while sacrificing brightness under standard marine conditions. The presence of these high-MGE strains in a coastal, temperate marine bacterium raises broader questions about what triggers MGE expansion and how these episodes impact host-microbe interactions.

Beyond variation in MGE content, *P. mandapamensis* from different hosts also differed in a set of accessory genes that may be linked to host or environmental conditions. In particular, low-MGE *S. roseigaster* strains shared a block of genes largely absent from other hosts, suggesting that host-associated populations may maintain distinct accessory gene repertoires. Although the functions of these genes were largely unknown, several were annotated as transport, motility, regulatory, and phage-related genes (Supp. Data 2). Although we cannot link these genes to specific phenotypes, their host-associated patterns provide testable hypotheses for future work on how environmental and host factors shape the accessory genome of *P. mandapamensis*. These findings demonstrate the value of investigating microbial variation at the strain level.

Our comparison of temperate and tropical *P. mandapamensis* strains provides insight into how environmental context can reshape symbiont genomes without sacrificing host specificity. Across hosts and habitats, *P. mandapamensis* retained a 14-gene locus that is absent from its close relative *P. leiognathi* and may contribute to the strong, species-level specificity observed in siphonfish light organs ^25,42^. Superimposed on this conserved core, we found divergence in various genes, indicating that selection acts on modular traits that tune luminescence, stress responses, and metabolism rather than on replacement of the symbiont. This pattern of a stable, specificity-conferring core with flexible strain-level genomic modules suggests that the *Siphamia*-*Photobacterium* partnership can accommodate ecological and functional variation while maintaining taxonomic fidelity. Future work on environmental monitoring, experimental cross-colonization, and transcriptomic profiling during symbiosis establishment and maintenance could clarify the extent of symbiont recruitment flexibility in siphonfish and its spatial distribution. These approaches can test whether hosts can acquire non-native strains when preferred partners are scarce and quantify the consequences of such shifts for symbiotic function. Understanding plasticity in the context of mutualisms is increasingly important for predicting how horizontally transferred symbioses respond to changing coastal environments in a rapidly changing ocean ^2,86^.

This study demonstrates that host-symbiont specificity in the *Siphamia*-*Photobacterium* symbiosis persists across temperate and tropical environments, despite *P. mandapamensis* exhibiting strain-level genomic and functional variation. Within this highly specific partnership, we found variation in luminescence-associated genes, IS expansion, host-associated accessory gene patterns, and the emergence of a non-luminous, host-derived strain. By studying symbiont diversity at the strain level, we show that specificity operates not only at the species level, but also between closely related host species that have overlapping ranges.

## METHODS

### Specimen Collection

Adult *Siphamia roseigaster* (n=3) and *Siphamia cephalotes* (n=11) were collected by SCUBA with a hand net from three sites near Sydney, NSW, Australia (Fig. 1; Supp. Data 1). Fish were euthanized according to IACUC-approved protocols (2021-12, IMSVBS), and light organs were dissected and placed into seawater-based agar medium (LSW-70; Lennox broth agar, 70% seawater). Agar tubes were shipped to the California Academy of Sciences (San Francisco, CA), where light organs were homogenized in 500 mL of sterile phosphate-buffered saline (PBS; Thermo Fisher) and plated in a dilution series on LSW-70 agar. We incubated plates at room temperature, selected 20-30 colonies, cultured the isolates in LSW-70 liquid medium, and cryopreserved them in 60% glycerol stock at -80°C for long-term storage. For downstream analyses, freezer stocks were revived, streaked onto fresh LSW-70 agar plates, and single colonies were inoculated into 3 mL of LSW-70 broth and grown overnight.

### Whole-genome sequencing

To complete long-read sequencing of the bacterial isolates, we extracted genomic DNA from cell pellets using the QIAGEN DNEasy Blood and Tissue Kit, following the manufacturer’s protocol. We performed ERIC-PCR (enterobacterial repetitive intergenic consensus PCR) fingerprinting using the DNA extractants and primers ERIC1R and ERIC2 (Supp. Methods ^87^). We visualized banding patterns using gel electrophoresis (1.5% agarose; GelDoc-It TS Imaging System), compared banding patterns, and selected unique isolates for sequencing. Genomic DNA from these representative isolates was re-extracted and purified using sparQ PureMag Beads (Quantabio; 0.5X bead-to-sample ratio). DNA concentrations were measured with the Qubit dsDNA HS Assay kit on a Qubit 3.0 fluorometer (Thermo Fisher) and standardized to 10 ng µl^−1^ for sequencing. We prepared sequence libraries with the Rapid (24) Barcoding Kit (Oxford Nanopore Technologies), pooled the libraries, and sequenced them on a MinION R10.4.1 flow cell.

### Pangenome and Statistical Analyses

After sequencing, we quality-filtered raw reads using Filtlong (v0.3.0; https://github.com/rrwick/Filtlong) to remove the lowest 5% by quality and any sequences shorter than 250 bp. Reads were assembled with Flye (v2.8.3)^88^, circularized using Circlator (v0.14.0)^89^, and polished with Medaka (v2.1.0; (https://github.com/nanoporetech/medaka) and Homopolish (v0.4.1)^90^ using ‘*Photobacterium*’ as the input genus. Assembly quality and completeness were assessed with BUSCO^91^ with the *Vibrionaceae* dataset, and potential contamination was screened with CheckM^92^. Genomes were scaffolded with RagTag (v2.1.0)^93^ and annotated with Prokka (v1.14.5)^94^ and Bakta^95^. Pangenome analysis was performed with Roary (v3.13.0)^96^ using a 95% identity threshold for core gene clusters. We created phylogenetic trees using IQ-TREE2 (v2.1.2)^97^ and visualized outputs using FigTree (v1.4.4) (http://tree.bio.ed.ac.uk/software/figtree/), Proksee^98^, and Phandango (v1.3.1)^99^. We identified mobile genetic elements and insertion sequences with MobileElementFinder (v1.1.2)^100^ and ISEscan (v1.7.3)^101^, while functional annotation was performed with eggNOG-mapper (v2.1.13)^102^. We used PlasmidVerify^103^ to identify plasmid replicon sequences. All programs were run using Conda v.25.3.1.

We generated pan- and core-cluster accumulation curves by permuting genome order 100 times within each host group (Sr, Sc, StP) and calculating the pangenome and core genome as genomes were added sequentially. We plotted the mean curves with ±1 SD across permutations. Using accessory genes from the Roary gene presence/absence matrix (core ≥ 95%), we calculated Jaccard distances between strains and performed a principal coordinates analysis on the resulting distance matrix (*cmdscale*). Kimura distances among IS copies were computed from multiple sequence alignments under a Kimura 2-parameter model, and distance distributions were visualized to identify bursts of IS expansion. To visualize the location of IS in the high-MGE strains, we built a consensus reference genome from an alignment of the eight high-MGE strains (Sr2 and Sc7.2). For each strain, ISEScan-predicted IS were projected onto the consensus via alignment-based coordinate conversion, and strain-specific tracks were annotated with ISEScan family assignments.

### Luminescence and Growth Rate Assays

We completed growth rate and luminescence assays under standard and variable salinity and temperature conditions using a multifunctional microplate reader (Varioskan ALF, Thermo Scientific, United States). From streaked plates, eight distinct isolates were selected as biological replicates, grown overnight in LSW-70 broth, and standardized to an OD_600_ of 0.25. Cultures were then diluted 1:20 into 96-well microplates containing LSW-70 adjusted to two salinity levels (5 and 26 ppt NaCl), with 26 ppt representing the standard condition. During each assay, plates were maintained at the target temperature (15°C, 25°C, or 30°C), and optical density (OD600) and luminescence were measured every 10 minutes for 24 hours, with pulsed shaking between measurements. To complete measurements for the 15°C condition, the plates were stored in a 15°C fridge, and manual measurements were taken every 30 minutes in 12-hour blocks over a 60-hour period. Media-only wells were included on each plate for blank-subtracting OD_600_ and luminescence values for downstream analyses. We summarized each replicate’s maximum growth rate as the slope of OD_600_ versus time during the exponential growth phase. Luminescence was reported as the maximum RLU achieved during the 24 h assay for each replicate and log_10_-transformed to reduce skew and stabilize residual variance. Growth and luminescence data were analyzed in R (v4.3.3) and GraphPad Prism (v10.5.0). Growth rate and luminescence were analyzed using linear mixed-effects models (*lme4* package)^104^ with host species, salinity, and temperature as fixed effects and strain as a random intercept. We initially used linear mixed-effects models with strain as a random intercept to account for unequal replication across strains, but the estimated variance for the strain effect was approximately zero (a singular fit), so we report results from the simpler linear model. For single-factor comparisons within individual assay conditions, we used one-way ANOVA and Tukey’s HSD tests with the *emmeans* package ^105^. To determine the emission profile of strains, we measured luminescence intensity in two wavelength channels (450 nm and 525 nm) on the Varioskan ALF for six biological replicates per strain, blank-subtracted using media-only wells, and calculated a 450/525 emission ratio for each replicate. Differences in 450/525 ratios among groups were compared using a one-way ANOVA followed by Tukey’s HSD post hoc comparisons (*emmeans*^105^).

## ACKNOWLEDGMENTS

We would like to acknowledge Paul V. Dunlap for his pioneering work on this system, and the Temple University presidential fellowship for partially supporting this work. We would like to thank Saniyah Johnson, Madelyn Akersten, Emma Román, Dr. Magdalena Warren, Matthew Soesanto, and Lily Hammer for their technical help and thoughts on this work.

## AUTHOR CONTRIBUTIONS

A.L. Gould and H.K. Osland conceived of the research. A. Gaisiner collected fish samples. H.K. Osland, E. Neff, and D. Hays carried out experiments. H.K. Osland carried out the analyses and wrote the initial manuscript. A.L. Gould funded the research and edited the manuscript.

## DATA AVAILABILITY

All raw sequence data (SRA) and genome assemblies (GenBank) generated in this study are publicly available at NCBI or will be released upon publication. The complete list of accession numbers is provided in Supplementary Table 1.

---

Supplemental Data 1. Collection metadata for siphonfish collection ^106,107^. Supplemental Data 2. Genes enriched in low-MGE *S. roseigaster* strains. Supplemental Methods. ERIC-PCR Conditions.

Supplemental Figure 1. (A) Growth rates of *P. mandapamensis* strains grouped by host species and MGE category under standard conditions (25°C, 26 ppt) (B) Growth rates under varying salinity and (C) temperature conditions.

Supplemental Figure 2. Growth and luminescence curves for representative *P. mandapamensis* strains at standard and varying salinity and temperature conditions.

Supplemental Figure 3. Phylogenetic tree of ISNCY.

Supplemental Figure 4. Phylogenetic tree of IS3.

Supplemental Figure 5. Phylogenetic tree of IS1182.

Supplemental Figure 6. Phylogenetic tree of IS630.

Supplemental Figure 7. Phylogenetic tree of IS5.

Supplemental Figure 8. Kimura distance plots of all high-MGE strains.

Supplemental Table 1. Assembly statistics and GenBank IDs of *P. mandapamensis* strains.

Supplemental Table 2. Number of singletons (unique genes) and plasmid replicon sequences per *P. mandapamensis* strain according to pangenome analysis.

Supplemental Table 3. Number of insertion sequences and composite transposons (MGEs) per *P. mandapamensis* strain (ISEScan and MEFinder).

Supplemental Table 4. Pairwise comparisons of growth rate and luminescence.

Supplemental Table 5. Strong candidate cargo genes based on putative composite transposon annotations by MobileGeneticElement Finder.

