## Supplemental for "High specificity meets genomic flexibility in the *Siphamia*-*Photobacterium* symbiosis"

**Supplemental Methods.** ERIC-PCR conditions.

ERIC-PCR was carried out using primer ERIC1R (5'- CACTTAGGGGTCCTCGAATGTA- 3') and ERIC2R (5'- AAGTAAGTGACTGGGGTGACGC- 3'). Reactions were performed in 25 µL volumes containing 50 ng genomic DNA, 2.5 µL of PCR buffer, 0.5 µL of dNTP, 1.0 µL of MgCl, 0.5 µLs of ERIC 1R, 0.5 µL of ERIC2R, and 0.3 µL of Taq. Amplification was performed with initial denaturation at 95°C for 7 min, followed by 35 cycles of denaturation at 90°C for 30s, annealing at 52°C for 1 min, and extension at 65°C for 8 min, with a final extension at 68°C for 16 min. PCR products were visualized on 1.5% agarose gels and visualized using a GelDoc-It TS imaging system.

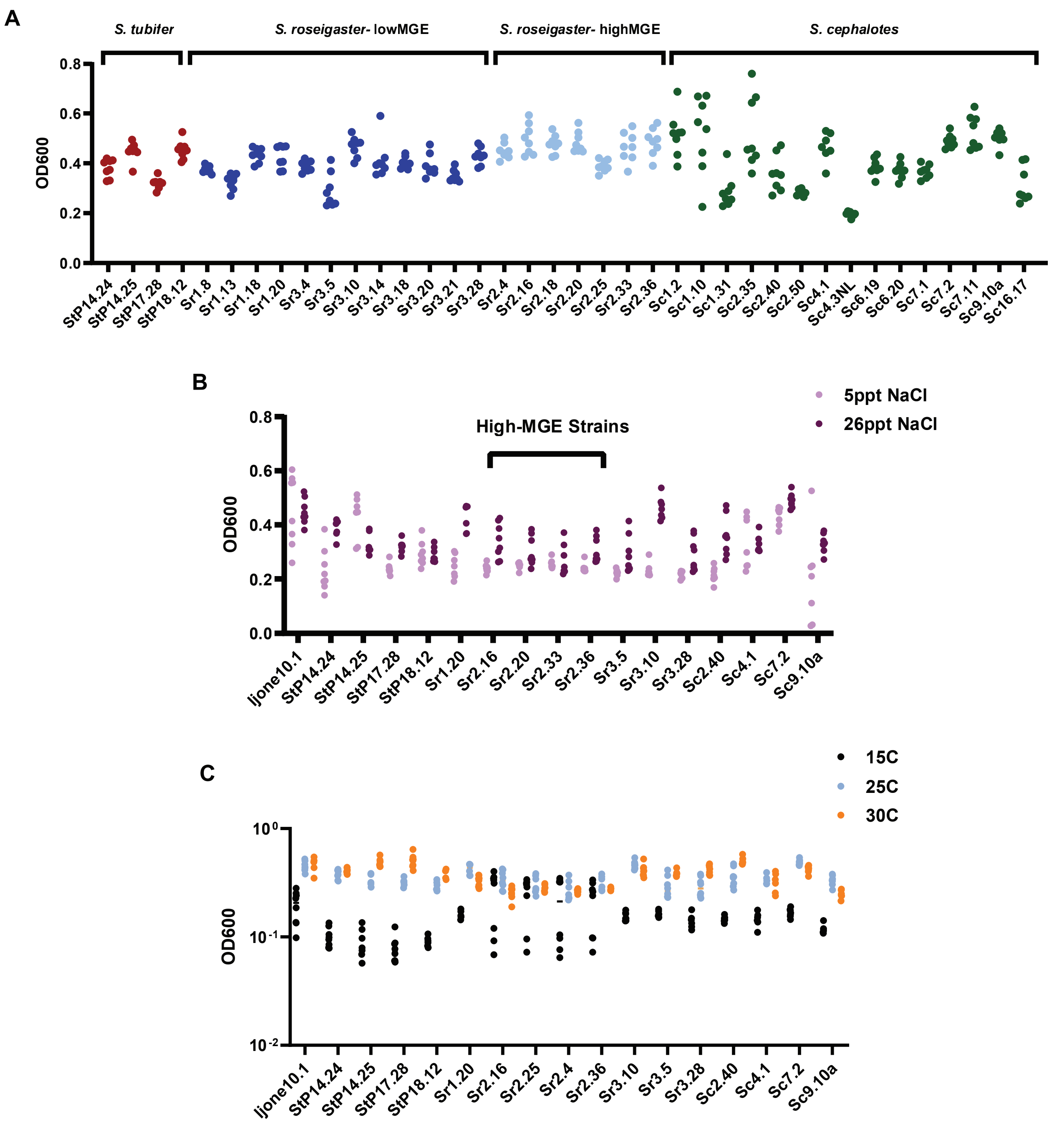


**Supplemental Figure 1.** (A) Growth rates of *P. mandapamensis* strains grouped by host species and MGE category under standard conditions (25°C, 26 ppt) (B) Growth rates under varying salinity and (C) temperature conditions.

**Supplemental Figure 2**. Growth and luminescence curves for representative *P. mandapamensis* strains at varying temperature conditions.

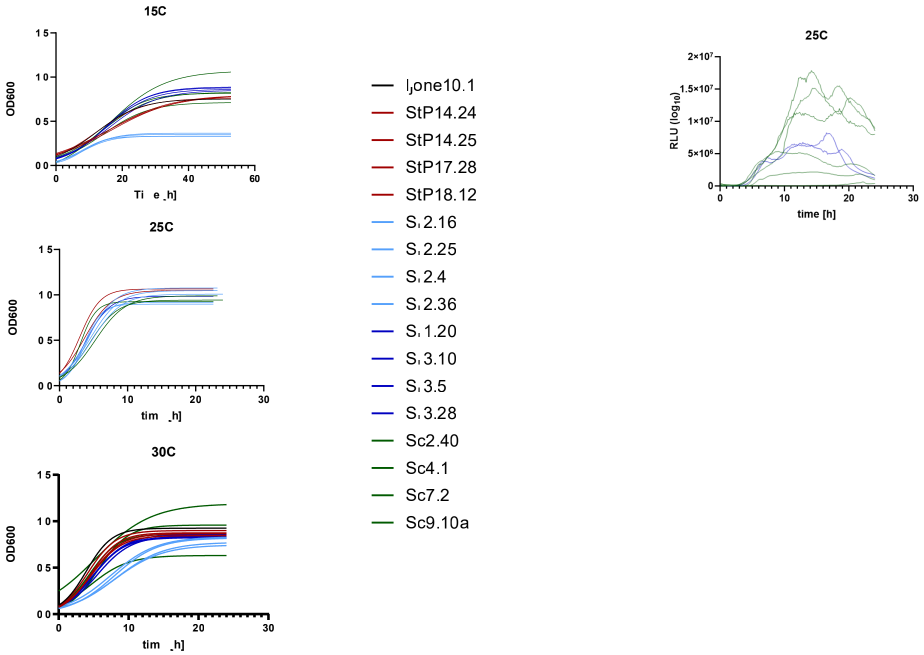

**Supplemental Figure 3.** Phylogenetic tree of ISNCY.

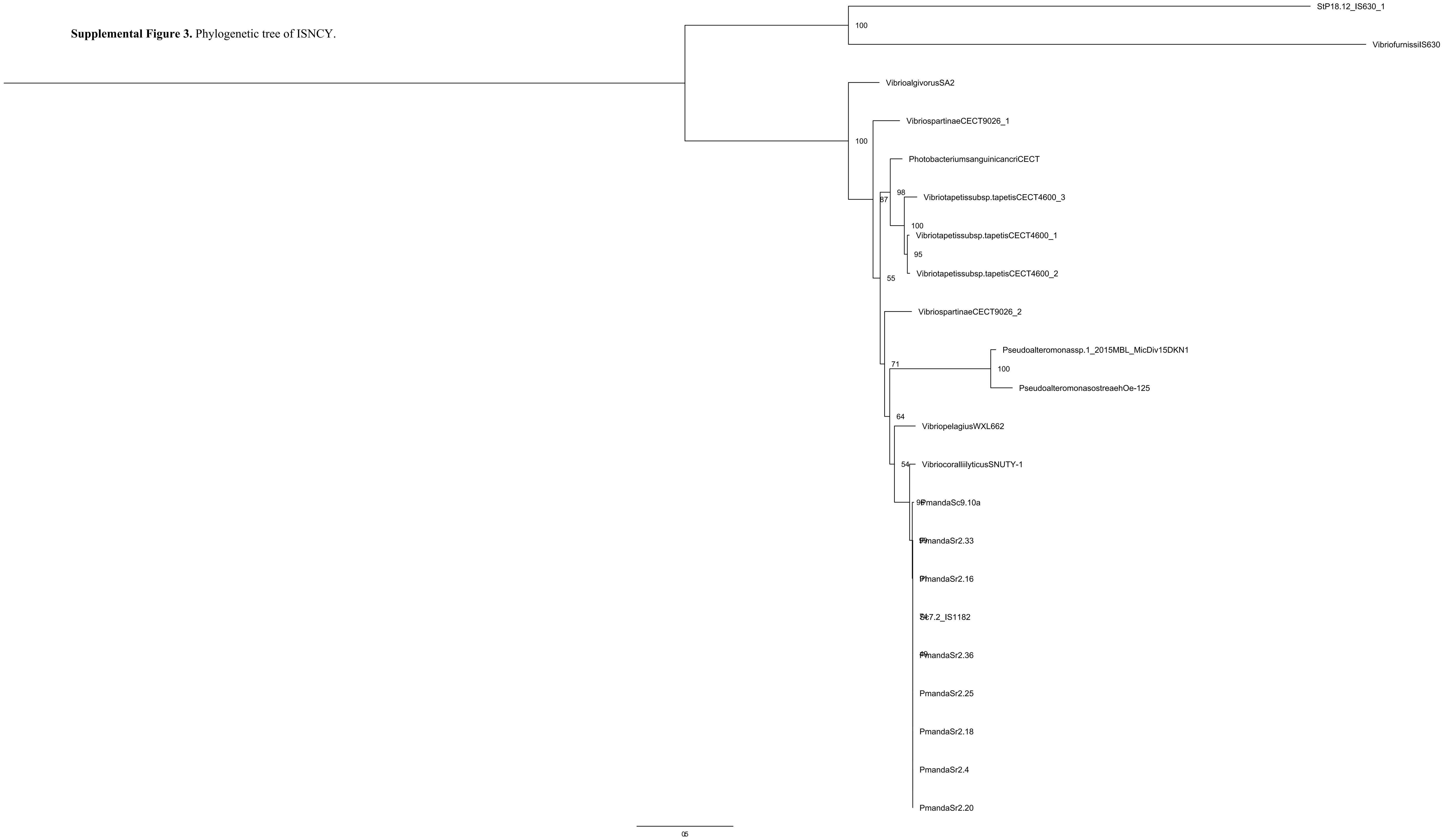

**Supplemental Figure 4.** Phylogenetic tree of IS3.

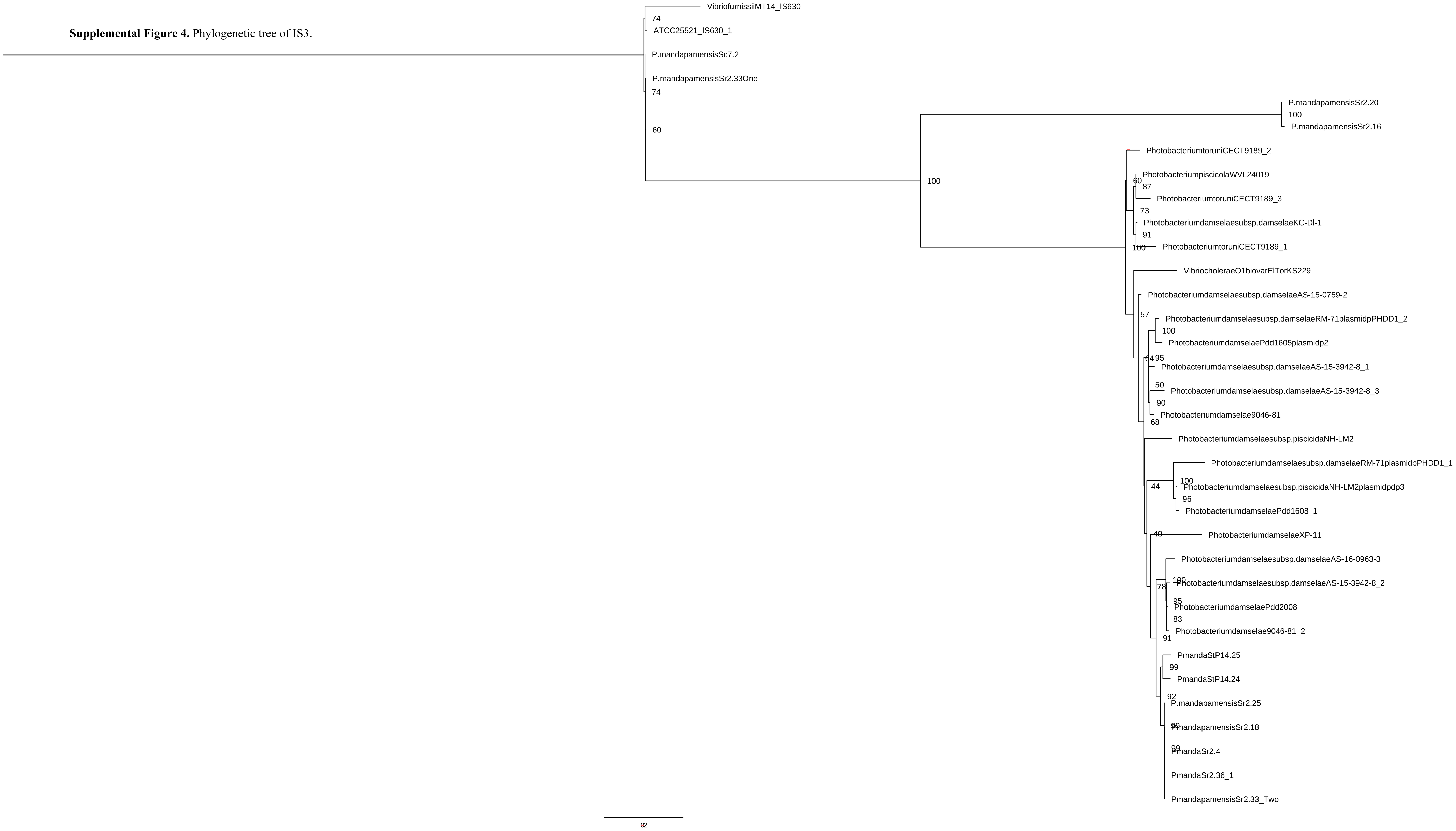

**Supplemental Figure 5.** Phylogenetic tree of IS1182.

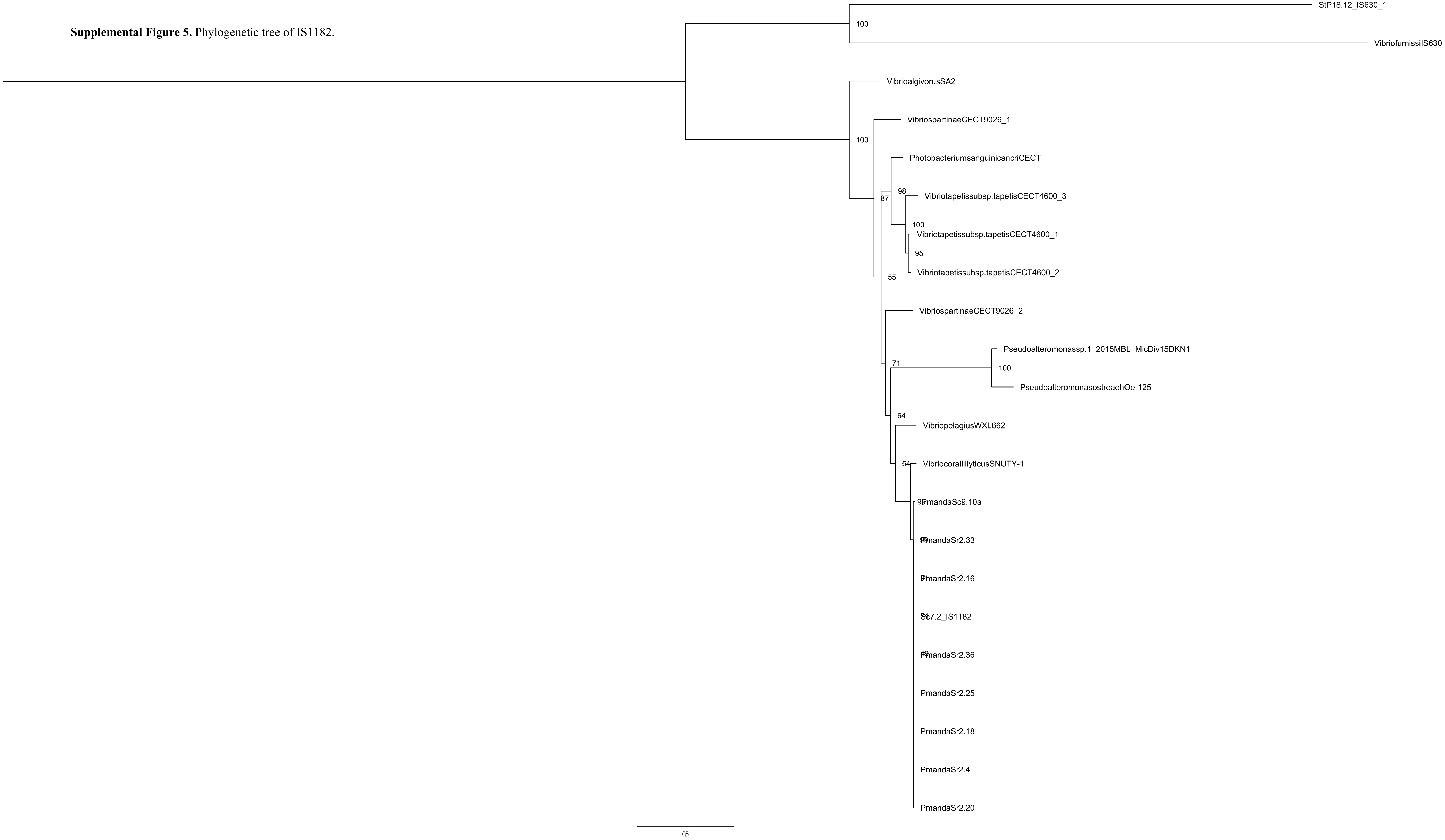

**Supplemental Figure 6.** Phylogenetic tree of IS630.

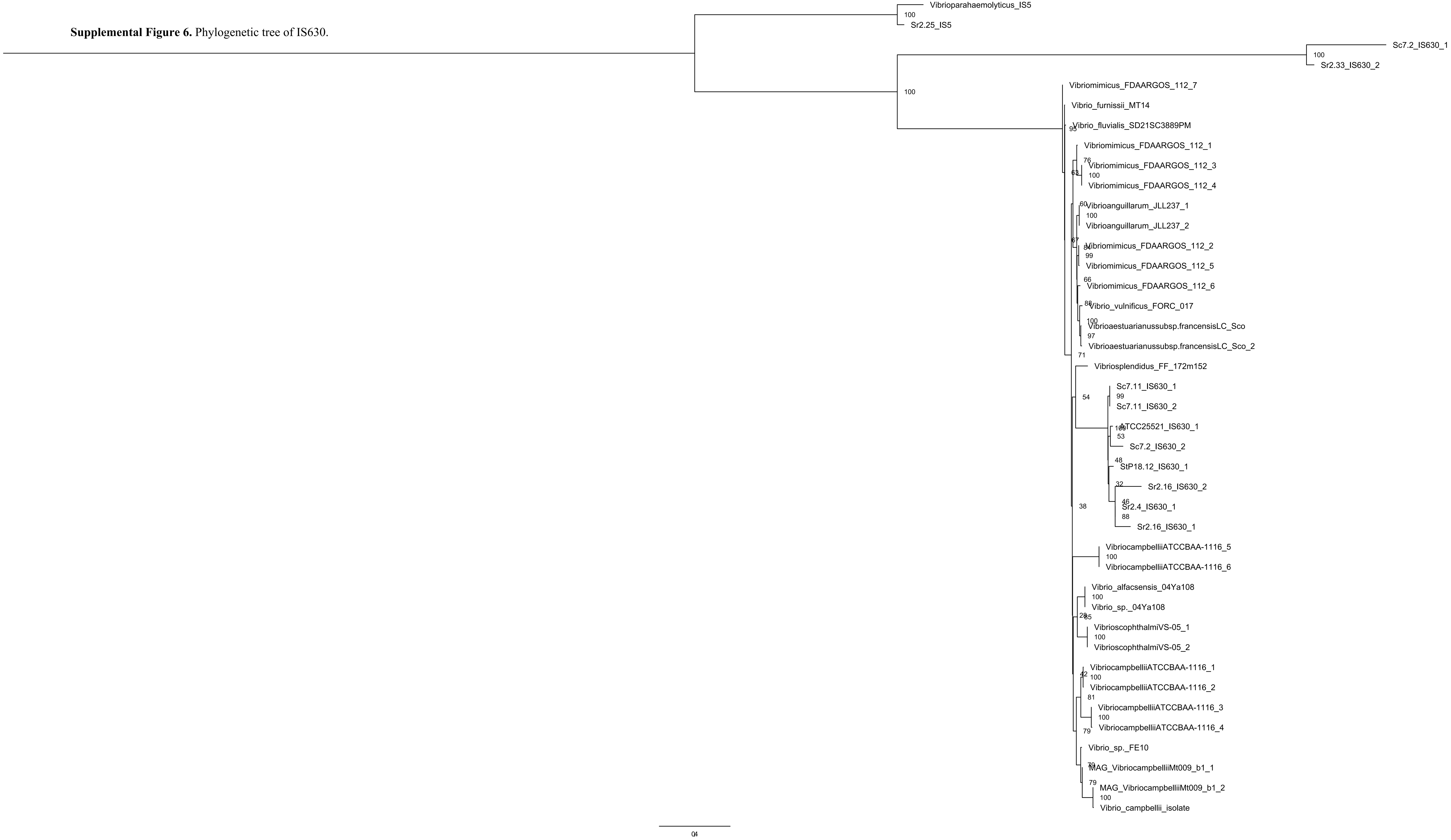

**Supplemental Figure 7.** Phylogenetic tree of IS5.

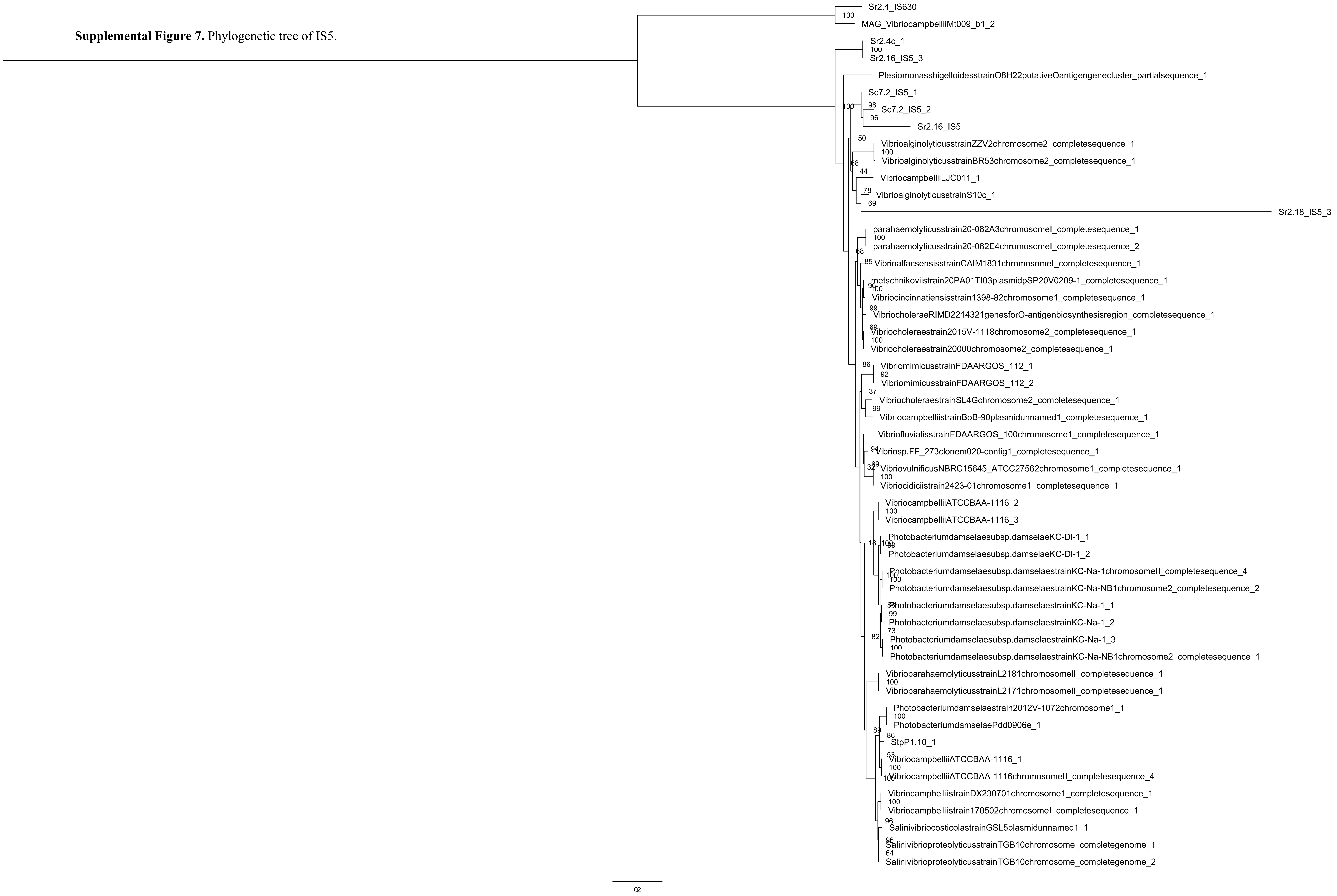

**Supplemental Figure 8.** Kimura distance plots of IS within high-MGE strains.

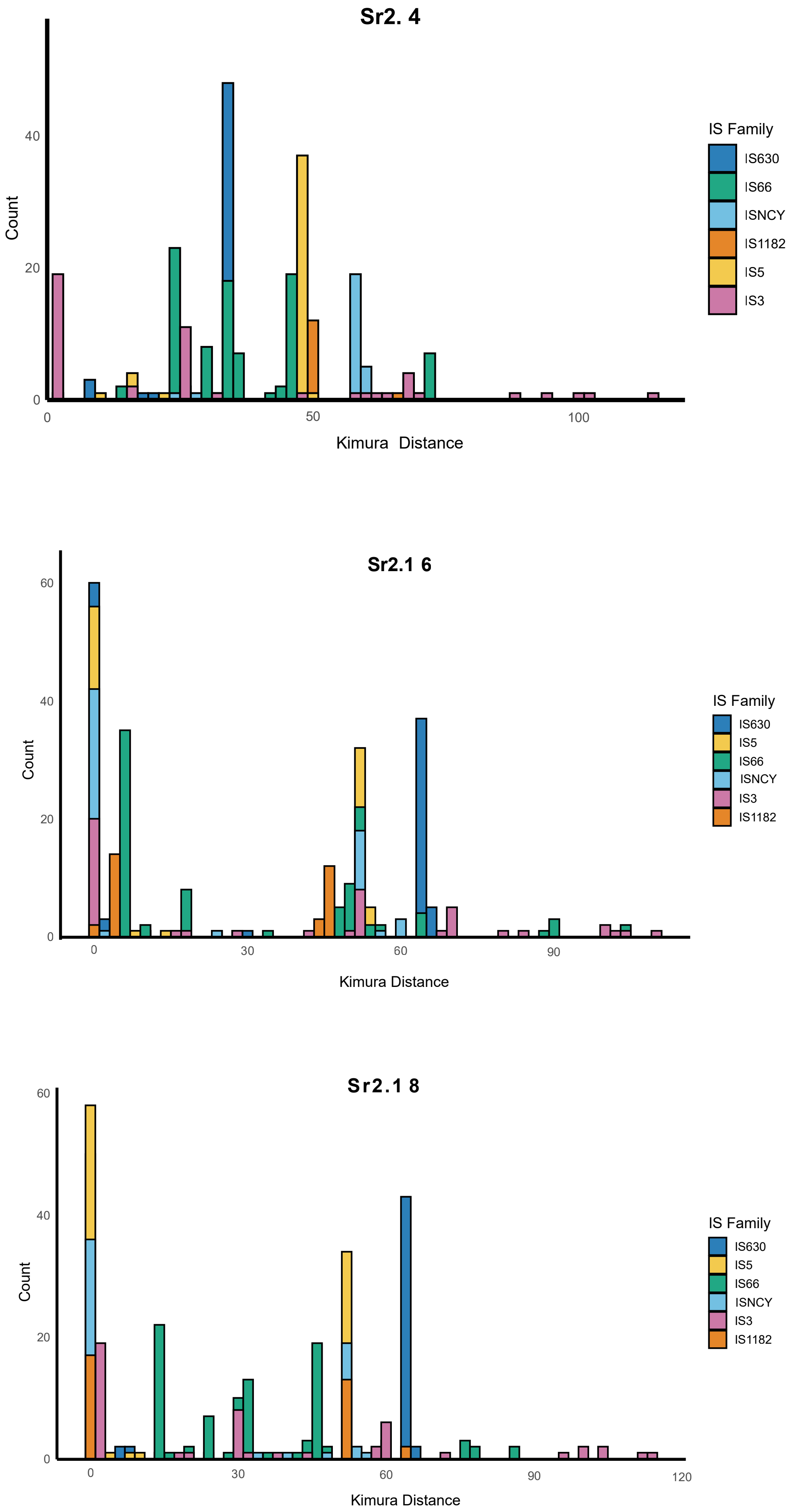

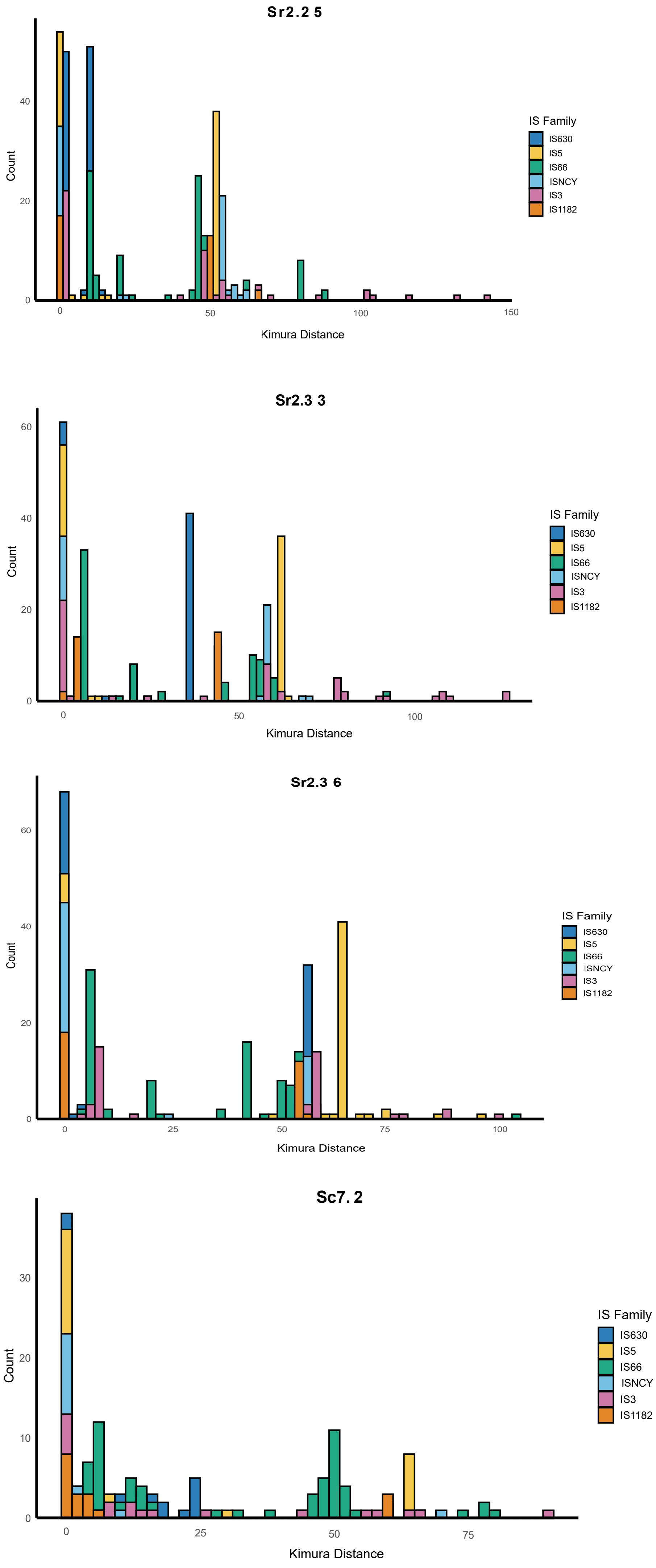

Sr2.2 5
Count
Kimura Distance
0
20
40
0
50
100
150
IS Family
IS630
IS5
IS66
ISNCY
IS3
IS1182
Sr2.3 3
Count
Kimura Distance
0
20
40
60
0
50
100
IS Family
IS630
IS5
IS66
ISNCY
IS3
IS1182
Sr2.3 6
Count
Kimura Distance
0
20
40
60
0
25
50
75
100
IS Family
IS630
IS5
IS66
ISNCY
IS3
IS1182
Sc7. 2
Count
Kimura Distance
0
10
20
30
0
25
50
75
IS Family
IS630
IS5
IS66
ISNCY
IS3
IS1182

**Supplemental Table 1.** Assembly statistics and GenBank IDs of *P. mandapamensis* strains.

| Strain ID | Total bp | Contigs | Largest contig | Mean Coverage (X) | N50 | GC (%) | BUSCO | checkM | Genbank ID |
|---|---|---|---|---|---|---|---|---|---|
| **lrivu 4.1** | 5268214 | 20 | 1730671 | 33 | 979827 | 40.98 | 95.9 | 0.27 | GCA_000509205.1 |
| **ATCC25521$^T$** | 4750881 | 3 | 3269131 | 18 | 3269131 | 41.02 | 99.2 | 1.27 | GCA_030685535.1 |
| **ljone10.1** | 5276714 | 8 | 3313411 | 55 | 3313411 | 41.34 | 100 | 1.11 | GCA_030716965.1 |
| **StP14.25** | 4763264 | 2 | 3202474 | 29 | 3202474 | 41.17 | 100 | 0.89 | Released upon publication |
| **Ik8.2** | 4783140 | 3 | 3218875 | 60 | 3218875 | 41.17 | 100 | 0.62 | GCA_030685395.1 |
| **StP18.31** | 4828605 | 3 | 3229357 | 76 | 3229357 | 41.18 | 100 | 0.59 | Released upon publication |
| **StP17.28** | 4740947 | 2 | 3230976 | 65 | 3230976 | 41.21 | 100 | 1.07 | Released upon publication |
| **StP18.12** | 4834338 | 3 | 3290233 | 22 | 3290233 | 41.09 | 100 | 0.59 | GCA_051046875.1 |
| **StP14.24** | 4745205 | 2 | 3187236 | 36 | 3187236 | 41.17 | 100 | 0.59 | GCA_051046865.1 |
| **Sc6.20** | 5019604 | 4 | 3200550 | 111 | 3200550 | 41.16 | 100 | 1.5 | Released upon publication |
| **Sc6.19** | 4982665 | 3 | 3195405 | 239 | 3195405 | 41.15 | 100 | 1.5 | Released upon publication |
| **Sc4.1** | 6066669 | 92 | 1591678 | 51 | 282935 | 41.09 | 100 | 4.31 | GCA_050084805.1 |
| **Sc1.31** | 4627104 | 11 | 2197283 | 60 | 2197283 | 43.61 | 100 | 2.98 | Released upon publication |
| **Sc2.49** | 4804769 | 13 | 3193716 | 36 | 3193716 | 41.2 | 100 | 1.03 | Released upon publication |
| **Sc1.2** | 4776751 | 14 | 2537776 | 23 | 2537776 | 41.15 | 99.2 | 1.27 | Released upon publication |
| **Sc9.10a** | 4729202 | 2 | 3246861 | 168 | 3246861 | 41.29 | 100 | 0.59 | GCA_048537455.1 |
| **Sc2.50** | 4723268 | 2 | 3224622 | 164 | 3224622 | 41.32 | 100 | 0.59 | GCA_049315015.1 |
| **Sc2.35** | 4717318 | 3 | 3213088 | 70 | 3213088 | 41.28 | 99.2 | 0.59 | Released upon publication |

| | | | | | | | | | |
|---|---|---|---|---|---|---|---|---|---|
| **Sc2.40** | 5324624 | 13 | 2785390 | 159 | 1498647 | 40.79 | 100 | 6.64 | Released upon publication |
| **Sc1.10** | 4880042 | 9 | 2785366 | 23 | 1498640 | 41.02 | 100 | 2.49 | Released upon publication |
| **Sc16.3** | 4771160 | 2 | 3236353 | 58 | 3236353 | 41.2 | 100 | 0.59 | GCA_048537465.1 |
| **Sc16.17** | 4799704 | 3 | 3235941 | 103 | 3235941 | 41.21 | 100 | 0.59 | Released upon publication |
| **Sc7.2** | 6867845 | 139 | 1254743 | 69 | 118364 | 40.98 | 100 | 11.49 | Released upon publication |
| **Sc7.1** | 4758435 | 5 | 2900434 | 27 | 2900434 | 41.19 | 100 | 1.41 | Released upon publication |
| **Sc7.11** | 4880562 | 3 | 3238926 | 110 | 3238926 | 41.23 | 100 | 0.86 | Released upon publication |
| **Sr2.33** | 5638392 | 17 | 3373868 | 56 | 3373868 | 41.5 | 100 | 0.81 | Released upon publication |
| **Sr2.20** | 5622561 | 13 | 3378511 | 52 | 3378511 | 41.51 | 100 | 0.81 | Released upon publication |
| **Sr2.16** | 5653832 | 22 | 3380506 | 82 | 3380506 | 41.49 | 100 | 1.09 | Released upon publication |
| **Sr2.36** | 5636714 | 12 | 3378510 | 137 | 3378510 | 41.49 | 100 | 0.81 | GCA_048537515.1 |
| **Sr2.25** | 5735811 | 10 | 3378535 | 54 | 3378535 | 41.45 | 100 | 0.81 | Released upon publication |
| **Sr2.4** | 5724430 | 11 | 2820637 | 83 | 2308030 | 41.47 | 100 | 0.91 | GCA_048537495.1 |
| **Sr2.18** | 5633965 | 12 | 3389354 | 36 | 3389354 | 41.47 | 100 | 1.45 | Released upon publication |
| **Sr3.28** | 5089675 | 4 | 3296021 | 119 | 3296021 | 41.2 | 98.4 | 0.91 | Released upon publication |
| **Sr3.20** | 5086295 | 6 | 3017087 | 164 | 3017087 | 41.19 | 100 | 0.91 | Released upon publication |
| **Sr1.20** | 5090337 | 5 | 2332925 | 101 | 1585241 | 41.23 | 94.3 | 0.91 | Released upon publication |
| **Sr3.18** | 5036172 | 8 | 3003917 | 54 | 3003917 | 41.13 | 99.2 | 1.11 | Released upon publication |
| **Sr3.10** | 5097880 | 4 | 3306367 | 83 | 3306367 | 41.22 | 100 | 0.91 | GCA_048537505.1 |

| | | | | | | | | | |
|---|---|---|---|---|---|---|---|---|---|
| **Sr3.5** | 5059785 | 6 | 3278724 | 47 | 3278724 | 41.19 | 100 | 0.91 | Released upon publication |
| **Sr1.18** | 5076394 | 4 | 3301000 | 172 | 3301000 | 41.23 | 100 | 0.91 | GCA_048537445.1 |
| **Sr3.4** | 5060449 | 6 | 3018813 | 82 | 3018813 | 41.15 | 97.6 | 1.04 | Released upon publication |
| **Sc4.3NL** | 5114211 | 11 | 3014476 | 16 | 3014476 | 41.14 | 98.4 | 0.98 | Released upon publication |
| **Sr1.7** | 5085481 | 4 | 3306355 | 30 | 3306355 | 41.19 | 100 | 0.64 | Released upon publication |
| **Sr1.11** | 5107651 | 6 | 3311960 | 76 | 3311960 | 41.11 | 100 | 2.92 | Released upon publication |
| **Sr3.21** | 5071752 | 4 | 3306358 | 67 | 3306358 | 41.21 | 100 | 0.91 | GCA_048537525.1 |
| **Sr1.8** | 5085241 | 5 | 3287580 | 23 | 3287580 | 41.14 | 100 | 1.18 | Released upon publication |
| **Sr3.14** | 5104964 | 4 | 3306338 | 74 | 3306338 | 41.19 | 100 | 0.91 | Released upon publication |
| **Sr1.13** | 5081840 | 4 | 3306384 | 56 | 3306384 | 41.2 | 99.2 | 0.91 | Released upon publication |

**Supplemental Table 2.** Number of singletons (unique genes) and plasmid replicon sequences per *P. mandapamensis* strain from pangenome analysis.

| Strain ID | Number of singletons | Number of plasmid replicon sequences |
|---|---|---|
| **lrivu 4.1** | 596 | - |
| **ATCC25521$^{T}$** | 342 | - |
| **ljone10.1** | 542 | - |
| **StP14.25** | 185 | - |
| **Ik8.2** | 140 | - |
| **StP18.31** | 167 | - |
| **StP17.28** | 113 | - |
| **StP18.12** | 290 | - |
| **StP14.24** | 161 | - |
| **Sc6.20** | 22 | 4 |
| **Sc6.19** | 27 | 3 |
| **Sc4.1** | 456 | - |
| **Sc1.31** | 260 | 11 |
| **Sc2.49** | 52 | 13 |
| **Sc1.2** | 105 | 14 |
| **Sc9.10a** | 120 | - |
| **Sc2.50** | 13 | - |
| **Sc2.35** | 17 | 3 |
| **Sc2.40** | 1 | 21 |
| **Sc1.10** | 5 | 21 |
| **Sc16.3** | 14 | - |
| **Sc16.17** | 35 | 3 |
| **Sc7.2** | 79 | 110 |
| **Sc7.1** | 33 | 5 |
| **Sc7.11** | 18 | - |
| **Sr2.33** | 93 | 17 |
| **Sr2.20** | 58 | 13 |
| **Sr2.16** | 68 | 22 |
| **Sr2.36** | 43 | - |
| **Sr2.25** | 137 | 10 |

| | | |
|---|---|---|
| **Sr2.4** | 106 | - |
| **Sr2.18** | 129 | 12 |
| **Sr3.28** | 6 | 4 |
| **Sr3.20** | 27 | 6 |
| **Sr1.20** | 8 | 5 |
| **Sr3.18** | 11 | 8 |
| **Sr3.10** | 8 | - |
| **Sr3.5** | 4 | 6 |
| **Sr1.18** | 7 | - |
| **Sr3.4** | 32 | 6 |
| **Sc4.3NL** | 534 | 11 |
| **Sr1.7** | 15 | 4 |
| **Sr1.11** | 67 | 6 |
| **Sr3.21** | 6 | - |
| **Sr1.8** | 29 | 5 |
| **Sr3.14** | 8 | 4 |
| **Sr1.13** | 5 | 4 |

**Supplemental Table 3.** Number of insertion sequences and composite transposons per *P. mandapamensis* strain (ISEScan and MEFinder).

| Strain ID | Number of IS (ISEScan) | Number of IS (MEFinder) | Number of Putative Composite Transposons (MEFinder) |
|---|---|---|---|
| **lrivu 4.1** | 61 | 14 | 7 |
| **ATCC25521$^T$** | 59 | 43 | 9 |
| **ljone10.1** | 127 | 79 | 37 |
| **StP14.25** | 5 | 1 | 0 |
| **Ik8.2** | 5 | 0 | 0 |
| **StP18.31** | 5 | 0 | 0 |
| **StP17.28** | 4 | 0 | 0 |
| **StP18.12** | 11 | 5 | 1 |
| **StP14.24** | 2 | 1 | 0 |
| **Sc6.20** | 40 | 20 | 5 |
| **Sc6.19** | 26 | 23 | 4 |
| **Sc4.1** | 180 | 132 | 65 |
| **Sc1.31** | 26 | 10 | 0 |
| **Sc2.49** | 15 | 8 | 0 |
| **Sc1.2** | 15 | 10 | 0 |
| **Sc9.10a** | 29 | 7 | 0 |
| **Sc2.50** | 12 | 3 | 0 |
| **Sc2.35** | 13 | 3 | 0 |
| **Sc2.40** | 28 | 7 | 0 |
| **Sc1.10** | 28 | 7 | 0 |
| **Sc16.3** | 15 | 6 | 0 |
| **Sc16.17** | 15 | 6 | 0 |
| **Sc7.2** | 320 | 233 | 95 |
| **Sc7.1** | 29 | 24 | 1 |
| **Sc7.11** | 30 | 14 | 1 |
| **Sr2.33** | 461 | 354 | 233 |
| **Sr2.20** | 463 | 357 | 236 |
| **Sr2.16** | 468 | 359 | 237 |
| **Sr2.36** | 465 | 350 | 241 |
| **Sr2.25** | 470 | 359 | 242 |

| | | | |
|---|---|---|---|
| **Sr2.4** | 461 | 362 | 257 |
| **Sr2.18** | 452 | 354 | 231 |
| **Sr3.28** | 9 | 2 | 0 |
| **Sr3.20** | 8 | 12 | 3 |
| **Sr1.20** | 8 | 2 | 0 |
| **Sr3.18** | 9 | 2 | 0 |
| **Sr3.10** | 9 | 2 | 0 |
| **Sr3.5** | 9 | 2 | 0 |
| **Sr1.18** | 8 | 2 | 0 |
| **Sr3.4** | 7 | 2 | 0 |
| **Sc4.3NL** | 7 | 2 | 0 |
| **Sr1.7** | 9 | 2 | 0 |
| **Sr1.11** | 8 | 2 | 0 |
| **Sr3.21** | 8 | 2 | 0 |
| **Sr1.8** | 11 | 4 | 0 |
| **Sr3.14** | 8 | 2 | 0 |
| **Sr1.13** | 8 | 2 | 0 |

**Supplemental Table 4.** Pairwise comparisons of growth rate and luminescence values among *P. mandapamensis* from different host siphonfish species based on linear mixed models. Estimates are differences in group means and Tukey-adjusted p-values. Luminescence values are $\log_{10}$-transformed.

| **Response** | **Contrast** | **df** | **Estimate** | **SE** | **t-value** | **p-value** |
|---|---|---|---|---|---|---|
| Growth rate | StP vs. Sc | 38 | -0.00697 | 0.044397 | -0.16 | 0.998598 |
| Growth rate | StP vs. Low-MGE Sr | 38 | 0.012374 | 0.045294 | 0.27 | 0.992755 |
| Growth rate | StP vs. High-MGE Sr2 | 38 | -0.06321 | 0.050075 | -1.26 | 0.59204 |
| Growth rate | Sc vs. Low-MGE Sr | 38 | 0.019343 | 0.028833 | 0.67 | 0.907432 |
| Growth rate | Sc vs. High-MGE Sr2 | 38 | -0.05624 | 0.035879 | -1.57 | 0.408875 |
| Growth rate | Low-MGE Sr vs. High-MGE Sr2 | 38 | -0.07558 | 0.036983 | -2.04 | 0.190331 |
| Luminescence ($\log_{10}$) | StP vs. Sc | 38 | -0.916 | 0.427 | -2.15 | 0.1598 |
| Luminescence ($\log_{10}$) | StP vs. Low-MGE Sr | 38 | -1.521 | 0.438 | -3.47 | 0.0074 |
| Luminescence ($\log_{10}$) | StP vs. High-MGE Sr2 | 38 | 0.751 | 0.476 | 1.58 | 0.4032 |
| Luminescence ($\log_{10}$) | Sc vs. Low-MGE Sr | 38 | -0.606 | 0.294 | -2.06 | 0.1866 |
| Luminescence ($\log_{10}$) | Sc vs. High-MGE Sr2 | 38 | 1.667 | 0.347 | 4.80 | 0.000176 |
| Luminescence ($\log_{10}$) | Low-MGE Sr vs. High-MGE Sr2 | 38 | 2.273 | 0.361 | 6.30 | 2.07E-06 |
